# Ligand-binding pockets in RNA, and where to find them

**DOI:** 10.1101/2025.03.13.643147

**Authors:** Seth D. Veenbaas, Jordan T. Koehn, Patrick S. Irving, Nicole N. Lama, Kevin M. Weeks

## Abstract

RNAs are critical regulators of gene expression, and their functions are often mediated by complex secondary and tertiary structures. Structured regions in RNA can selectively interact with small molecules – via well-defined ligand binding pockets – to modulate the regulatory repertoire of an RNA. The broad potential to modulate biological function intentionally via RNA-ligand interactions remains unrealized, however, due to challenges in identifying compact RNA motifs with the ability to bind ligands with good physicochemical properties (often termed drug-like). Here, we devise *fpocketR*, a computational strategy that accurately detects pockets capable of binding drug-like ligands in RNA structures. Remarkably few, roughly 50, of such pockets have ever been visualized. We experimentally confirmed the ligandability of novel pockets detected with *fpocketR* using a fragment-based approach introduced here, Frag-MaP, that detects ligand-binding sites in cells. Analysis of pockets detected by *fpocketR* and validated by Frag-MaP reveals dozens of newly identified sites able to bind drug-like ligands, supports a model for RNA secondary structural motifs able to bind quality ligands, and creates a broad framework for understanding the RNA ligand-ome.

## INTRODUCTION

A subset of RNAs fold into complex secondary and tertiary structures capable of forming ligand-binding pockets that can engage small molecules with high affinity and specificity. RNA molecules lie upstream of most biological functions, both encoding proteins and broadly regulating gene expression. Small molecules that bind to and modulate the function of RNA thus have the potential to regulate diverse processes, including the levels of otherwise “undruggable” protein targets lacking well-defined ligand binding pockets (1–3). Further, non-coding RNAs present a vast scope of unexploited targets to manipulate biological processes (4). However, the promise of selectively targeting RNA with small molecules remains incompletely realized. Our current molecular understanding of RNA-ligand interactions is limited to a few classes of highly structured RNAs, including riboswitches (5), ribozymes (6), and ribosomal RNAs (rRNAs) (7). The lack of an understanding of the types of RNA structures that form high-affinity and selective interactions with small-molecule ligands remains a major challenge for ligand discovery efforts targeting RNA.

Pockets in RNA have previously been computationally detected using approaches originally developed for finding pockets in proteins, including V3 (8, 9) and mkgrid (10, 11), which are rolling probe-based approaches, and PocketFinder (12, 13), which is energy based. In addition, the ROBIN database uses machine learning to characterize RNA-binding ligands (14) and SHAMAN leverages RNA-focused molecular dynamics and fragment docking to identify small-molecule binding sites (15). These studies suggest that ligand-binding pockets in RNA and protein molecules have similar bulk properties of volume, buriedness, and solvent accessibility (9, 11, 13, 14). In contrast, RNA pockets have been proposed to be less hydrophobic and to bind more rod-like ligands (9, 11, 13, 16), features which are generally viewed unfavorably for medicinal chemistry (17). Prior work tended to include (nearly) all known ligand-bound RNA structures – containing a preponderance of simple organic molecules and less-complex stem-loop RNA structures – in contrast to focusing on RNA structures that bind to ligands with favorable (often called drug-like) physicochemical properties.

There is also intense debate regarding what kinds of specific RNA structures are capable of harboring pockets able to bind ligands with favorable physicochemical properties. The current dialogue centers on the relative importance of targeting simple stem-loop or bulge containing motifs, which may have stronger current biological validation, versus targeting complex RNA motifs which can potentially form more selective interactions with small molecules, but are harder to identify (3, 18–20). Most recent work has focused on the former class of simple RNA targets.

We posited, first, that differences in physicochemical properties between proteins and RNAs require that pocket-finding algorithms be optimized *specifically* for RNA structures. Second, pockets that bind ligands with favorable physicochemical properties (and are plausibly drug-like) should be prioritized to best understand RNA-ligand interactions in the context of structure-based ligand discovery (3, 21).

Here we introduce *fpocketR* and Frag-MaP, two frameworks for identifying ligandable pockets in RNA. *fpocketR* is a software package optimized to identify, visualize, and characterize pockets in RNA, built around the open-source pocket detection software *fpocket* (22). We optimized *fpocketR*, focusing on the limited examples of RNAs with known complex structures that bind small-molecule ligands with favorable physicochemical properties. *fpocketR* can be used as a discovery tool to identify novel pockets that specifically bind drug-like ligands in both apo (ligand free) and (low resolution) dynamic RNA tertiary structures. *fpocketR* provides users the flexibility to analyze single RNA structures or RNA ensembles, including experimentally solved structures, molecular dynamics simulations, or from computational modeling. We experimentally validated the ligandability of pockets detected by *fpocketR* using Frag-MaP. Frag-MaP uses photocrosslinkable small-molecule probes and mutational profiling (MaP) technology (23, 24) to identify the nucleotide position of RNA-ligand interactions in a direct experimental step, in cells. Finally, we analyzed pocket-forming RNA secondary structures identified using *fpocketR* and propose a framework for understanding RNA secondary structures that specifically interact with favorable (drug-like) small-molecule ligands.

## RESULTS

### Geometry-based analysis reveals differences in protein versus RNA pockets

We used the open-source, geometry-based pocket finding framework, *fpocket*, to detect ligandable pockets in RNA. *fpocket* places alpha spheres throughout the structure of a biomolecule, where every alpha sphere is in contact with the center point of exactly four atoms in the biomolecule (**Fig. 1A**). The radius of each alpha sphere reflects the local curvature of the biomolecule (22), allowing solvent accessible cavities to be identified as groups of appropriately sized alpha spheres. Ligand-binding pockets are detected by clustering groups of alpha spheres that are close in three-dimensional space. Finally, pockets are characterized and scored based on their physical and electrostatic properties. The default *fpocket* algorithm is widely used and successful for detecting ligandable pockets in proteins (22, 25).

**Figure 1.**
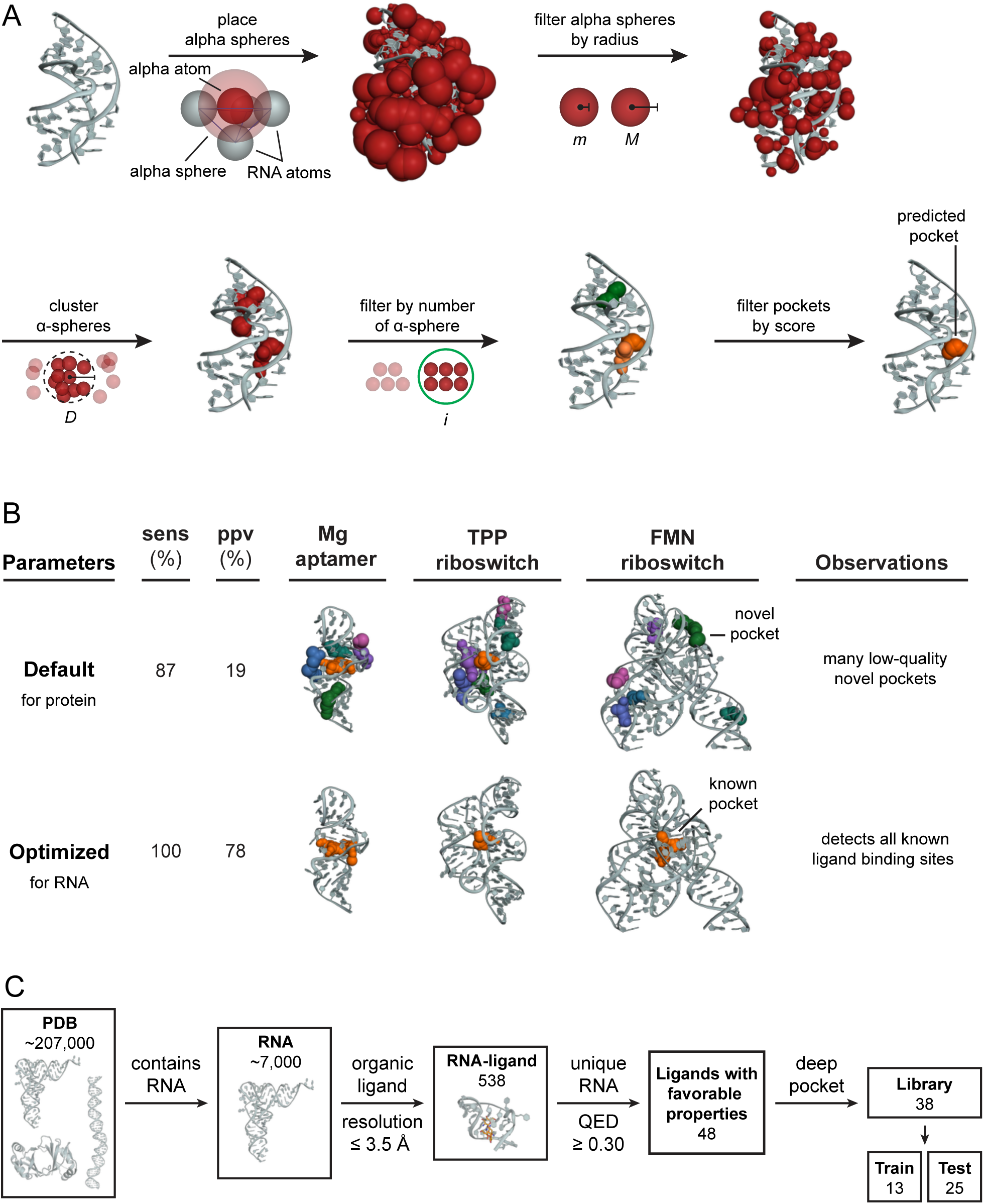
Geometry-based pocket finding approach and optimization for detecting sites in RNAs that bind ligands with favorable physicochemical properties. (*A*) Illustration of the *fpocket* algorithm and use of alpha spheres to detect pockets (PDB: 1F27) (47). (*B*) *fpocket* performance before and after optimization using a curated library of RNAs in complex with high QED ligands. Distinct pockets are indicated by differently colored alpha spheres. (Examples shown, PDB: 1Q8N, 2GDI, 5KX9) (48–50). (*C*) Curation and abundances of drug-like RNA-ligand complexes in the Protein Data Bank. RNA-ligand complexes were filtered from the HARIBOSS database (11). QED quantifies eight fundamental properties of a small molecule ligand into a single metric of drug-likeness (21).

The *fpocke*t algorithm was optimized to detect pockets in proteins and tuned for the properties of amino acids (**Fig. 1A**). When *fpocket* is applied to protein structures, the highest scoring pocket corresponds to the known ligand-binding site 83% of the time (22). We applied *fpocket* to a curated library of RNA-ligand complexes, selected for inclusion of drug-like ligands. Using the default, protein-optimized, parameters *fpocket* detects many (incorrect) pockets for each RNA structure, and the highest scoring pocket overlaps with the known ligand-binding site only 63% of the time. More worryingly, many pockets detected using the default parameters are located on the RNA surface or in non-selective and solvent-exposed grooves, making these “pockets” poor targets for selective, small-molecule ligands (**Fig. 1B**). The discrepancy in *fpocket* performance between protein and RNA structures, first, reveals that a refinement of the algorithm is required to reliably detect RNA pockets capable of binding ligands with favorable physicochemical properties and, second, emphasizes there are fundamental differences between the ligand-facing surfaces of RNA and protein.

### A curated structure database of drug-like ligands bound to RNA

To optimize *fpocket* for RNA, we curated a library of short (≤ 200 nt), high resolution (≤ 3.5 Å) RNA-ligand complexes from the 538 structures available in the non-redundant version of the HARIBOSS database (**Fig. 1C**) (11). We selected non-redundant RNAs containing ligands with favorable physicochemical properties, defined as a quantitative estimate of drug-likeness (QED) score (21) of ≥ 0.3. The QED score is an evolution of Lipinski’s rules that ranks small molecules based on eight fundamental physicochemical properties including molecular mass, hydrophobicity, number of rotatable bonds, and number of hydrogen bond donors and acceptors. With the QED metric, overall favorable molecular properties can compensate for one or more less desirable features to provide a holistic view of ligand quality in a single value that ranges from 0 (worse) to 1 (best). Requiring a QED score of ≥ 0.3 naturally excludes ligands that bind to or intercalate with RNA non- or semi-specifically.

In the following text, for simplicity, we refer to ligands with QED (drug-likeness) scores ≥ 0.3 as drug-like RNA-binding ligands. However, we note that few of the small molecules discussed here, or known to bind RNA generally, are true drugs. Instead, we use the term drug-like to emphasize compounds with good physicochemical properties, consistent with general medicinal chemistry convention. By these definitions, we identified 48 high-resolution structures of RNA-ligand interactions in which the small molecule ligand is plausibly drug-like (**Fig. 1C**). The paucity of structures in this list emphasizes that we have much to learn about the ligandability of RNA.

### *fpocketR*: *fpocket* optimized for RNA

Of the 48 RNA-ligand complexes, 38 had deep, well-defined pockets. We divided these 38 complexes into a training set (*n* = 13), used for optimization of *fpocketR* (**Table S1**), and a test set (*n* = 25), for validation (**Fig. 1C**, **Table S2**). Four core parameters (*m, M*, *D, i*) in the *fpocket* algorithm strongly affect RNA pocket detection by controlling alpha sphere size and clustering (**Fig. 1A**). We systematically evaluated 1820 combinations of these four parameter values in a three round multi-objective optimization to maximize sensitivity (sens) and positive predictive value (ppv) for detecting the known ligand-binding sites in our RNA-ligand training set (**Fig. S1, Table S3**).

Optimization of *fpocket* yielded parameters that accurately detect binding sites for drug-like ligands in RNA (**Fig. 1B** and **S1**). We incorporated these optimized parameters into *fpocketR*, a custom python package that detects, characterizes, and visualizes pockets in RNA. The optimized parameters (*m* = 3.0, *M* = 5.7, *D* = 1.65, and *i* = 42) used in *fpocketR* differ markedly from the default *fpocket* parameters (*m* = 3.4, *M* = 6.2, *D* = 2.4, and *i* = 15). Critical changes were to require both smaller alpha spheres and larger numbers of alpha spheres to comprise a pocket. Using these changes, the probability that the top-scoring pocket overlapped with a known ligand binding site increased from 63% (*fpocket*) to 92% (*fpocketR*), corresponding to a five-fold reduction in false positives (**Table S4**). Furthermore, *fpocketR* identified 100% of known ligand-binding sites in the training and test sets with a minimum positive predictive value of 78% (**Fig. 1B** and **2**). *fpocketR* also detected eight novel pockets that do not overlap known ligand-binding sites. These novel pockets appear to be high quality (buried, sufficient volume, supporting π-π stacking) (**Fig. 2**) and may bind cryptic ligands (suggesting the true positive predictive value is notably higher than 78%). For example, we detected a novel pocket in the SAM-III riboswitch, buried in the RNA structure between a multi-helix junction and a bulge, that is less solvent exposed than the known ligand-binding site (**Fig. S2**). *fpocketR* significantly outperformed *fpocket* in detecting binding sites for drug-like ligands in RNA structures (**Fig. 2B**; **Table S4**).

**Figure 2.**
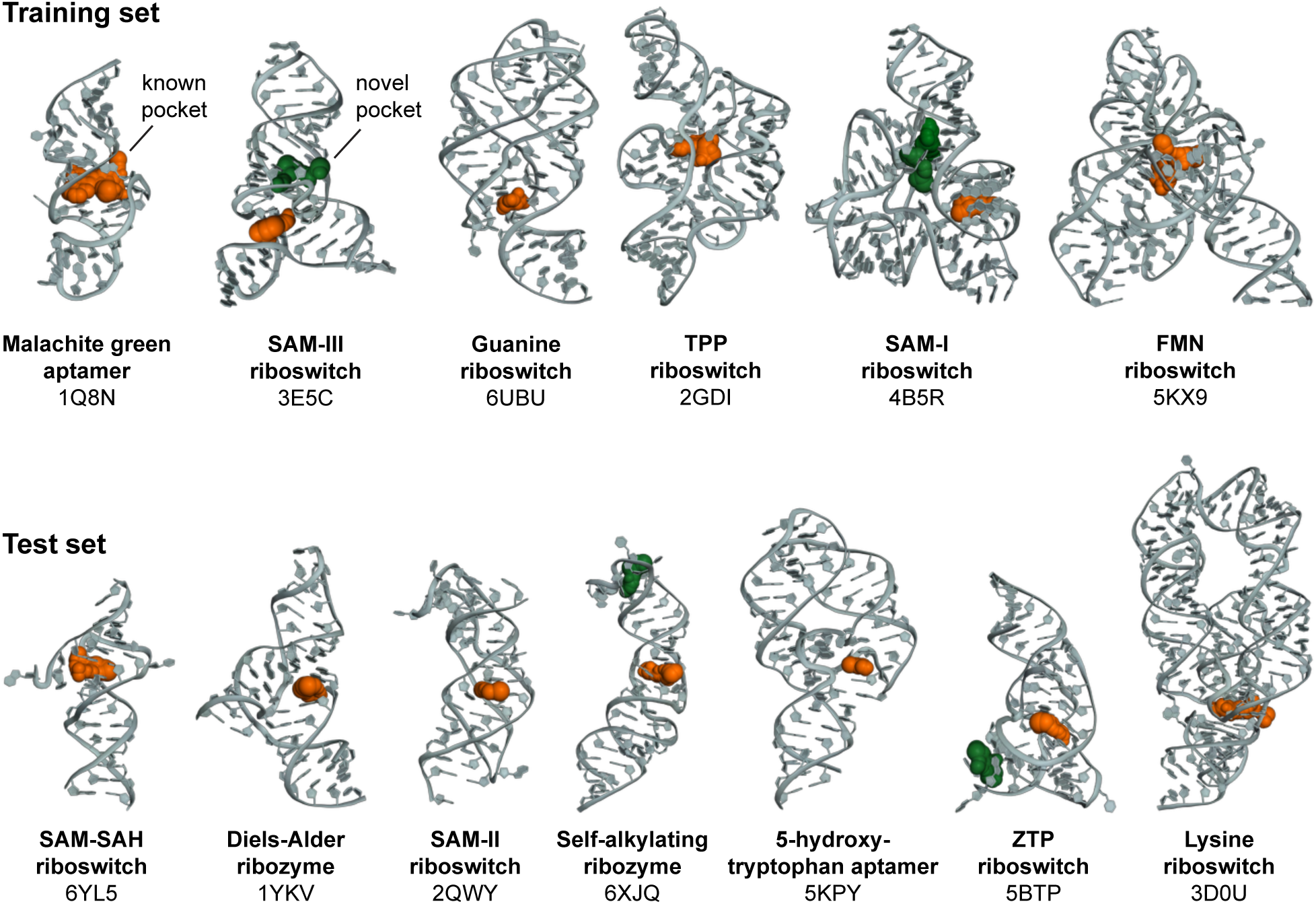
Pockets detected in RNA structures with *fpocketR*. Selected examples from the training and test sets are shown. Known and novel pockets indicated in orange and green, respectively.

### Frag-MaP: Experimental validation of *fpocketR* predictions

We applied *fpocketR* to the *E. coli* (PDB: 7K00) and *B. subtilis* (PDB: 7AS8) ribosomes (26, 27) and identified 46 and 52 pockets in the 23S rRNAs, respectively (**Fig. S3**). The ribosomal pocket-ome identified in this analysis includes most known ligand-binding sites for drug-like ligands, including binding sites for the antibiotics linezolid, sparsomycin, and spectinomycin. However, most predicted pockets are novel, suggesting the universe of RNA pockets capable of binding drug-like ligands might be quite large.

We therefore experimentally evaluated if the novel pockets identified by *fpocketR* bind drug-like small molecules in cells. We developed Frag-MaP, which leverages tri-functionalized probes containing a small-molecule fragment (MW ≤ 300) linked to a photocrosslinkable diazirine and “clickable” alkyne handle (**Fig. 3A**) (28, 29). Probes penetrate cells and crosslink to nucleotides proximal to their RNA-ligand binding sites, yielding RNA-ligand adducts. Frag-MaP exploits mutational profiling (MaP) (23, 24) for the step that identifies ligand-binding sites. MaP is direct and experimentally concise, and yields low sequencing and library-preparation biases, high sensitivity, and single nucleotide resolution. RNA-ligand adducts induce mutations in the cDNA synthesized during reverse transcription, which are quantified through massively parallel sequencing. These improvements are analogous to the advantages of using MaP to read out chemical probing experiments (23, 24). Frag-MaP sites are identified as nucleotides that interact with a fragment probe with significantly higher mutation rates compared to a control probe bearing a simple methyl group (**Fig. 3B**).

**Figure 3.**
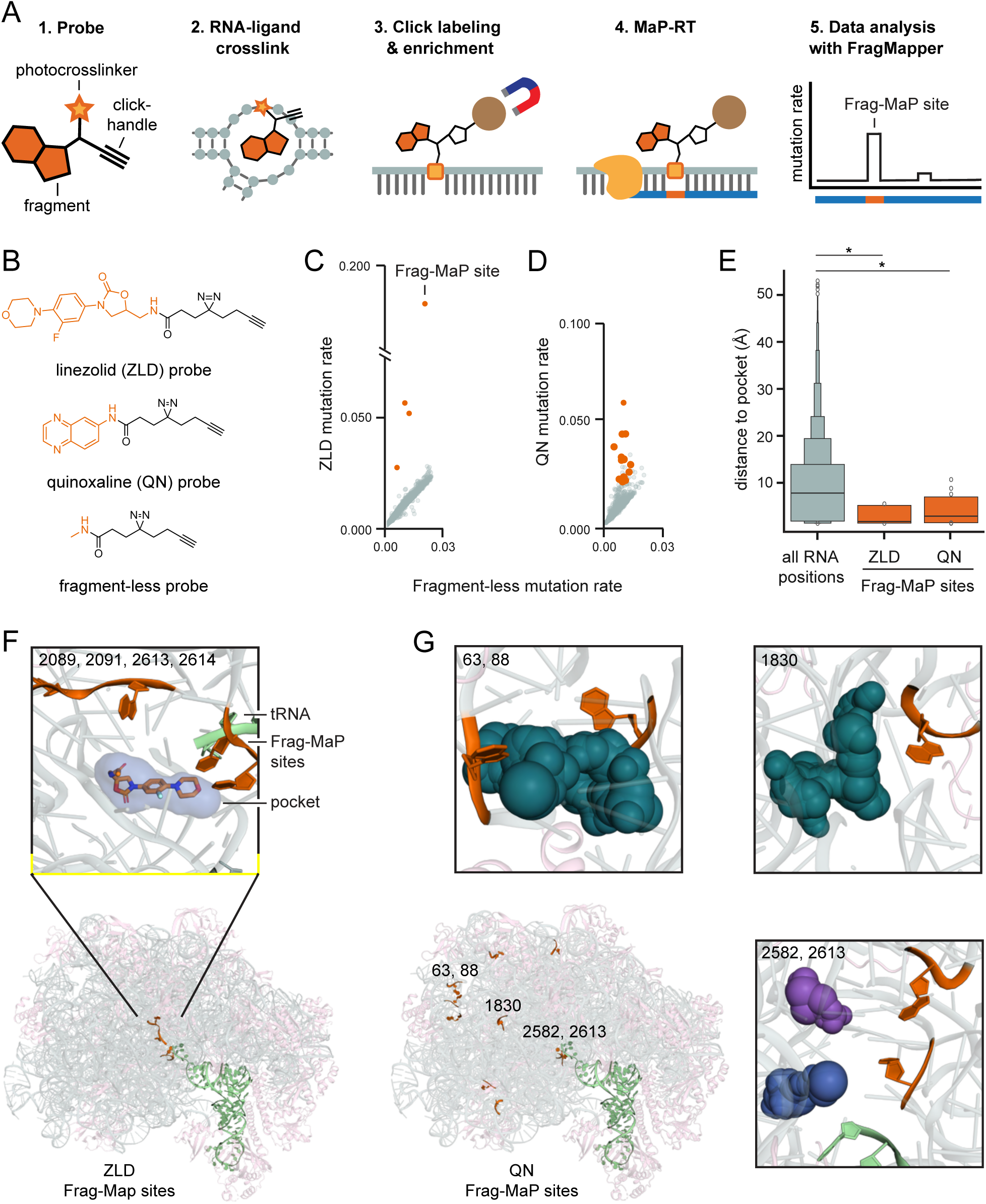
Experimental detection of RNA-ligand binding sites by Frag-MaP. (*A*) Frag-MaP experimental scheme. (*B*) Structure of high-QED probes used in Frag-MaP experiments. Mutation rate of Frag-MaP sites (orange, labeled) detected in the 23S rRNA for (*C*) linezolid and (*D*) quinoxaline probes. (*E*) Distance to a pocket predicted by *fpocketR* from all nucleotides in the 23S rRNA compared to Frag-MaP sites for linezolid and quinoxaline. *, p < 0.03; Mann Whitney test. Note: Frag-MaP sites occur at residues with relative solvent accessible surface area (rSASA) spanning 0.05 to 0.89 (0.0 and 1.0 equal to buried versus fully accessible). rSASA is not significantly different for Frag-MaP sites (mean: 0.44, stdev: 0.25) compared to all RNA residues (mean: 0.43, stdev: 0.22); p = 0.52, Mann-Whitney test. (*F*) Visualization of all Frag-MaP sites detected using the linezolid-based probe. Frag-MaP sites are orange; linezolid is shown bound in the peptidyl transferase center (PDB: 7AS8) (27). (*G*) Visualization of the most reactive Frag-MaP sites detected in the 23S rRNA using the quinoxaline-based probe. Pockets detected using *fpocketR* are shown as spheres.

We first validated the Frag-MaP strategy using a known RNA-ligand interaction based on a functionalized linezolid probe (**Fig. 3B**). Linezolid is a fragment- and drug-like (QED = 0.79) antibiotic with activity against Gram-positive bacteria and binds with low micromolar affinity to a pocket in the peptidyl transferase center (PTC) in the 23S rRNA (30, 31). The pocket in which linezolid binds is robustly predicted by *fpocketR* (**Fig. 3F**). The acetamidomethyl group is oriented toward the exit tunnel and provides a vector to functionalize linezolid without interfering with binding. We treated *B. subtilis* cells with the linezolid probe and detected four Frag-MaP sites as positions highly reactive toward the linezolid probe (mutation rates: 3-18%) relative to the fragment-less control probe (median mutation rate: 0.5%) (**Fig. 3C, Fig. S4A**). The four Frag-MaP sites detected for the linezolid probe (nucleotides 2089, 2091, 2613, 2614) are hundreds of nucleotides apart in primary sequence but lie within 7 Å of the linezolid binding site in three-dimensional space (**Fig. 3F, Fig.S4A**). Frag-MaP thus detects RNA-ligand binding sites with nucleotide resolution in cells, with high sensitivity and specificity.

We next screened for new ligand-binding sites in the *B. subtilis* 23S rRNA using 6- aminoquinoxaline (QED = 0.57) (**Fig. 3B**) and fragment 206 (QED = 0.81) probes (**Fig. S3C**). The quinoxaline probe yielded ten Frag-MaP sites with mutation rates up to 6% (**Fig. 3D**). Some of these sites were tens of nucleotides apart from each other in sequence, but clustered at compact locations in the 23S rRNA (**Fig. 3G**). Strikingly, 9 out of 10 Frag-MaP sites occurred at pockets predicted by *fpocketR* with a median distance of 4.5 Å (**Fig. 3E, Fig. S4B**). Probe 206 yielded an additional 4 Frag-MaP sites, 3 of which are near pockets (median distance of 5 Å) (**Fig. S4C**). Both probes bound to pockets in functionally interesting areas within the large ribosomal subunit. The quinoxaline probe bound to a pocket formed by a pseudoknot within the exit tunnel (nucleotides: 63 and 88) and to three nucleotides in the L1 stalk (nucleotides: 2145, 2154, 2198). Both the quinoxaline (nucleotides: 2582, 2613) and 206 (nucleotides: 2613, 2614) probes also bound at the P-site within the PTC, adjacent to the linezolid binding site. Identification of these site, in functionally important regions, demonstrate the ligandability and relevance of pockets detected by *fpocketR*.

Most (89%) ligand binding sites detected by Frag-MaP are close in three-dimensional space (median distance 4.8 Å) to pockets detected by *fpocketR.* Frag-MaP sites are significantly closer to pockets identified by *fpocketR* than expected by chance and occur at residues with both high and low solvent accessibility (**Fig. 3E**). The juxtaposition of pockets – predicted by *fpocketR* and validated experimentally by Frag-MaP – emphasizes that pockets detected by *fpocketR* are reflective of authentic, in-cell ligand-binding sites for drug-like small-molecule ligands.

### Pocket ligandability: validation of sites for drug-like small molecules

In a second validation strategy, we evaluated 17 RNA-ligand complexes comprised of antibiotics bound to bacterial rRNAs to assess the capacity of pockets detected by *fpocketR* to bind drug-like ligands. The 17 small-molecule ligands bind the 30S and 50S rRNAs at ten distinct binding sites, span multiple chemical classes, and have QED scores ranging from 0.01 (not drug-like) to 0.89 (highly drug-like) (**Fig. 4A, Fig. S5**). This is a rigorous test of *fpocketR* as rRNAs are significantly larger than and not evolved to bind small-molecule ligands compared to the RNA aptamers and riboswitches used to train *fpocketR*.

**Figure 4.**
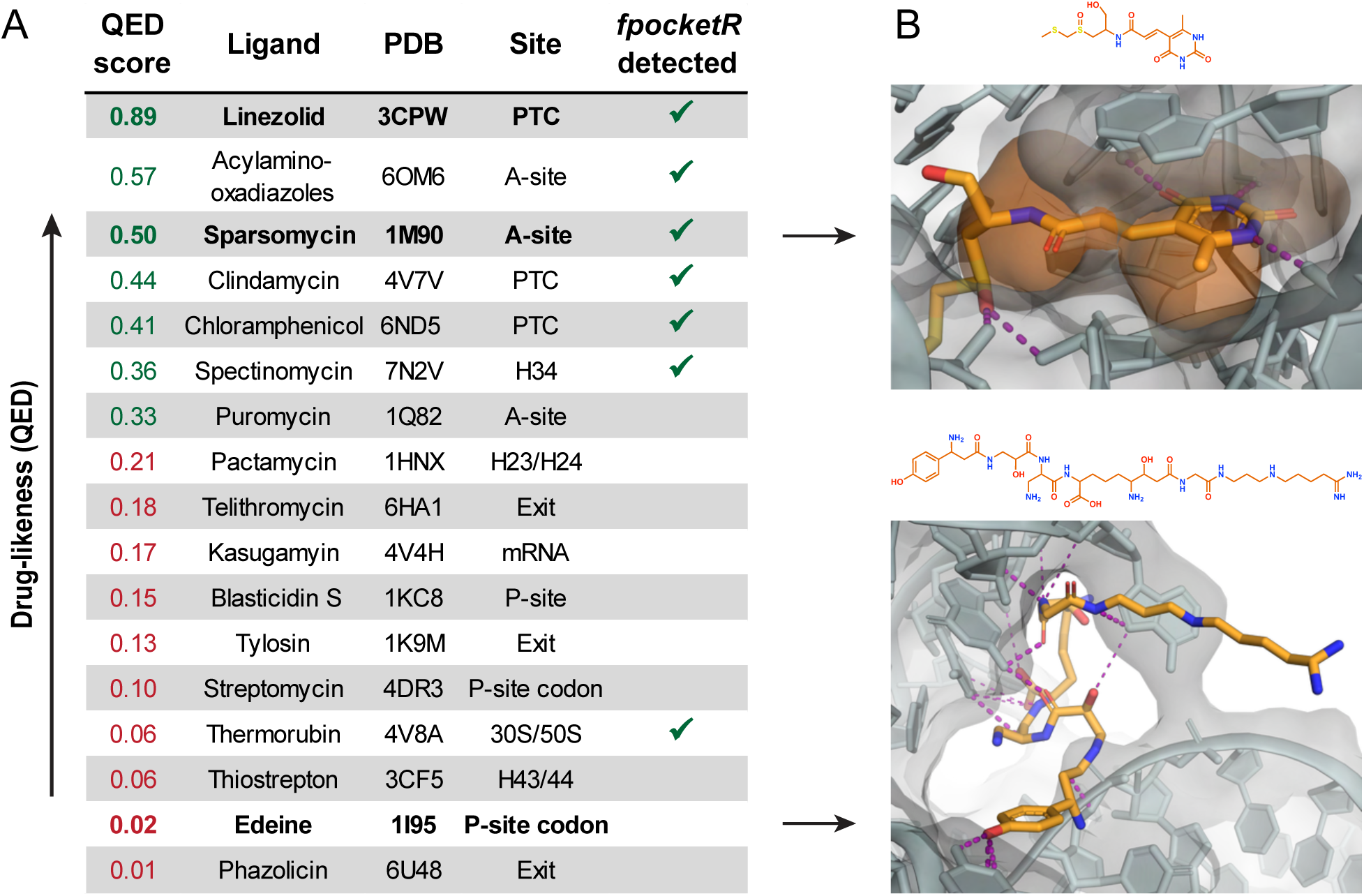
Pockets detected by *fpocketR* in the bacterial rRNA, compared to small-molecule antibiotic binding sites. (*A*) Ligand binding sites, listed by (QED) drug-likeness score. Sites detected with *fpocketR* indicated with check marks. (*B*) Contrasting examples of a drug-like ligand (sparsomycin) that binds a well-defined pocket (orange) and a non-drug-like ligand (edeine) that binds a non-pocket-like structure. An illustration of ligands bound near the PTC is provided in **Fig. S5**.

The known ligand-binding site was detected as a pocket by *fpocketR* for 7 of the 17 RNA-ligand complexes, across 4 distinct binding sites. *fpocketR* differentiated ligand-binding sites based on the drug-likeness of the bound ligands. *fpocketR* detected six of the seven pockets for ligands with a QED score above the training threshold of 0.3 (**Fig. 4A**). In addition, *fpocketR* detected the pocket for the highly conjugated ligand thermorubin that forms extensive π-π interactions with rRNA at the junction of the 50S and 30S ribosomal subunits. This pocket appears able to bind drug-like ligands, even though the ligand in this case is not drug-like.

There are clear structural differences between ligand-binding sites that interact with druglike and non-druglike ligands (**Fig. 4B**). Pockets that bind drug-like ligands are buried, generally encapsulate their ligands, and form both hydrogen bonding and π-π interactions. By contrast, RNA motifs that interact with less drug-like ligands, such as aminoglycosides and the extended edeine chain, are characterized by a high number of hydrogen bonding interactions and are located at solvent exposed RNA surfaces (**Fig. 4B**). All low-QED (and thus undrug-like) ligands in our analysis are natural products. Apparently, bacteria can evolve anti-ribosome antibiotics that do not follow human-compiled rules for drug-likeness and, further, can target ribosome regions that do not contain conventional pockets.

In sum, based on both direct experimental interrogation with Frag-MaP (**Fig. 3**) and by evaluating drug-like and non-drug-like molecules that bind the ribosome (**Fig. 4**), we validate that *fpocketR* selectively identifies pockets in RNA capable of binding ligands with drug-like physicochemical properties. We now use *fpocketR* to understand features of ligand-binding pockets in RNA.

### Detection of pockets in unliganded (apo) structures

We used *fpocketR* to examine binding sites in both apo (ligand-free) and dynamic structures. We analyzed paired ligand-free (apo) and ligand-bound (holo) RNA structures for ten RNA-ligand complexes spanning a wide-range of QED scores (0.23 to 0.89; **Table S5**). Most of these structures were determined by X-ray crystallography, so crystal packing partially stabilized the apo state. *fpocketR* detected the known ligand binding site in 9 of the 10 holo complexes, only failing to detect the ligand binding site for the least drug-like ligand, phosphoribosyl pyrophosphate (QED 0.23). *fpocketR* also detected the known ligand binding site in 9 of 10 apo (ligand-free) structures, including RNAs with global ligand-induced conformation changes (for example, the TPP riboswitch). The size, shape, and position of predicted pockets are often affected by ligand-induced conformational changes in the RNA structure, meaning that pockets in apo structures do not always perfectly align with known ligand bind-sites (**Fig. 5A**). However, *fpocketR*-detected pockets in the apo structures reliably determined the local capacity of an RNA to bind a small-molecule ligand.

**Figure 5.**
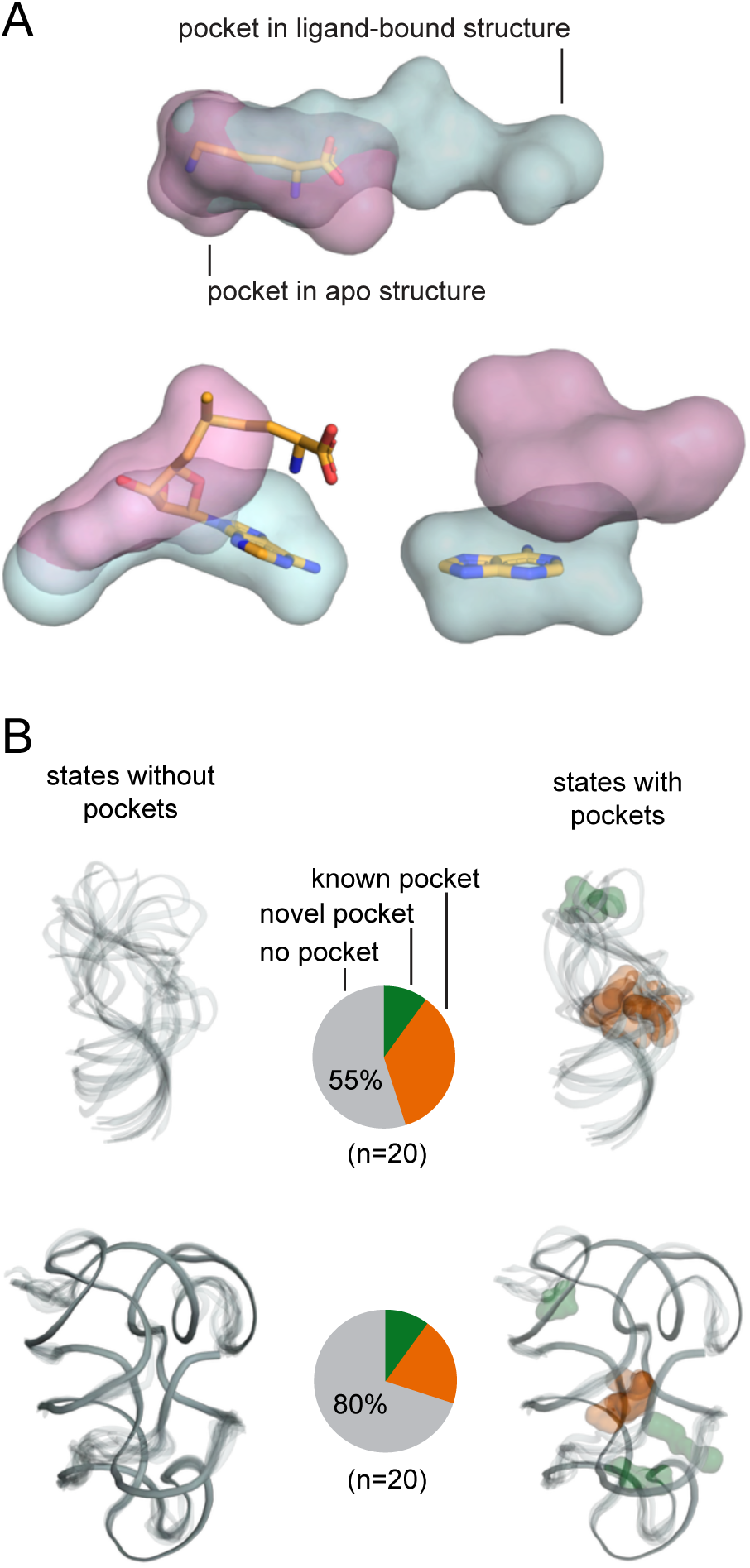
Detection of ligand-binding sites in apo (ligand-free) sites and in RNA ensembles. (*A*) Pockets detected by *fpocketR* in apo (pink) and ligand-bound (blue) RNA structures for the lysine riboswitch, SAM-I riboswitch, and adenine riboswitch (PDB: 3D0U & 3D0X, 3IQN & 3IQP, 5SWE & 5E54) (51–53). (*B*) Pockets detected in individual RNA states for ensembles of HIV-TAR, measured by NMR (PDB: 7JU1) (33), and the SAM-IV riboswitch, measured by cryo-EM (PDB: 6WLQ) (34). Known and novel pockets, detected by *fpocketR*, are orange and green, respectively. Structures are depicted as transparent backbones; pockets are shown as transparent surfaces.

RNA molecules often populate structural ensembles and the underlying structural dynamics influence biological function and response to ligand binding (32). This intrinsic dynamism in RNA structure also means that RNA structures in ensembles, especially for smaller RNAs, are visualized at modest resolution. We visualized pockets in a carefully refined NMR-based model of the HIV-1 TAR RNA hairpin (33). Of the 20 structures modeled for the TAR RNA, seven (35%) contained pockets overlapping the binding site for the known ligand (**Fig. 5B**). Similarly, an ensemble of structures was reported for the apo state of the of the SAM-IV riboswitch (determined by cryo-EM; 3.7 Å resolution) (34). In this case, pockets formed in four states (20%) (**Fig. 5B**). Novel pockets were observed for a single state for both RNAs. Thus, in these two examples of unliganded RNA structures with multiple states, pockets formed in only a minor subset of conformations but were concentrated at known ligand-binding sites. Small molecule binding then selects for the subset of structures with a ligandable pocket. Importantly, *fpocketR* finds ligandable sites even in ensembles based on modest-resolution structure models.

### Tripling the known universe of RNA pockets able to bind drug-like ligands

Non-redundant, high-resolution structures for RNAs are limited, and are especially sparse for RNAs bound to ligands with good (drug-like) physicochemical properties (**Fig. 1C**). Prior to this study, roughly 50 pockets capable of binding drug-like ligands had ever been visualized, these are those found in small RNAs (**Fig. 1C** and **Fig. 2)** and in the ribosomal RNAs (**Fig. 4**). Based on *fpocketR*, this work substantially expands this universe to 138, and could be made larger by extending the analysis developed here to additional RNAs.

### RNA secondary structures that harbor pockets

We examined RNA structures from the training set, test set, bacterial rRNAs, and a group II intron and identified all nucleotides that form pockets detected by *fpocketR*, and thus likely to bind drug-like ligands. We then annotated the nucleotides that form pockets in secondary structure diagrams (**Fig. 6**). In larger multi-domain RNAs, such as the ribosome or group II intron, pockets form both from nucleotides localized in secondary structure space (local interactions) and from nucleotides that become juxtaposed via through-space interactions involving multiple domains (long-range interactions) (**Fig. 7**, **Fig. S6-S7**, **Tables S6-S9**).

**Figure 6.**
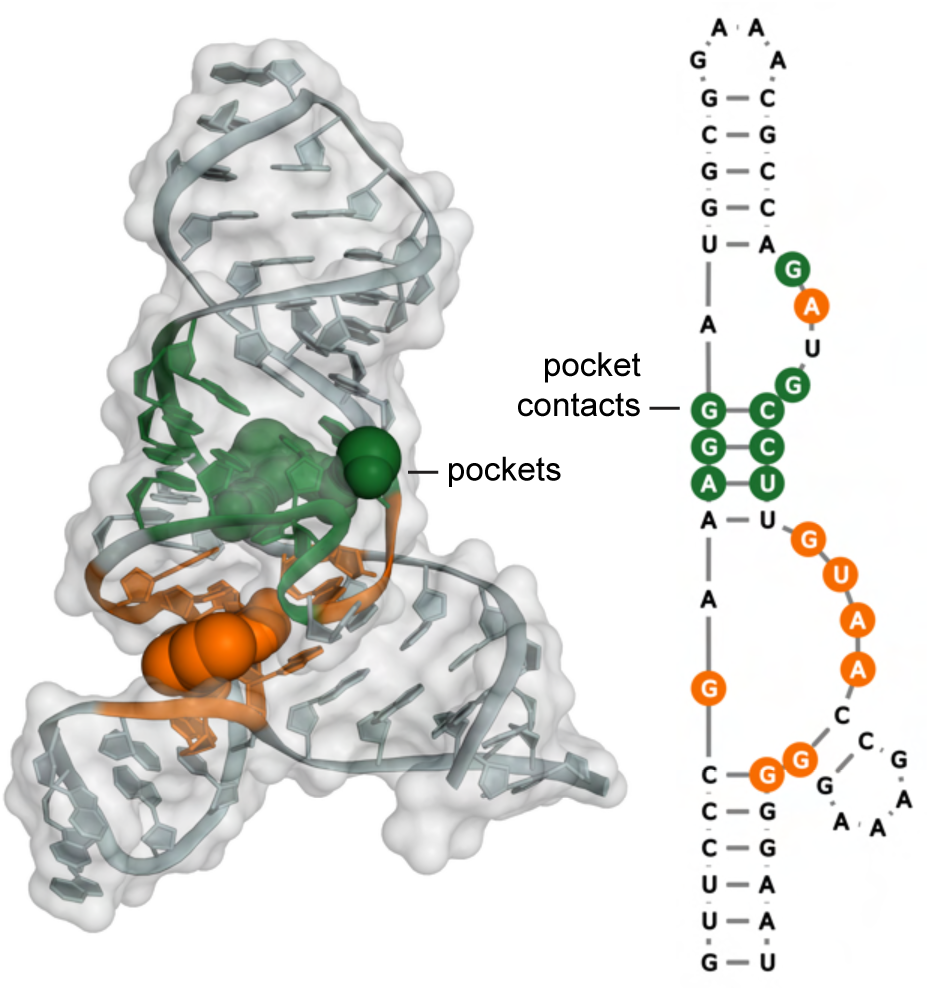
Scheme for mapping RNA pockets, visualized in three-dimensional structures, onto secondary structure diagrams. Nucleotides are colored to match the pocket they contact, in both tertiary and secondary structure images (PDB: 3E5C) (54). In this structure, both pockets are local.

**Figure 7.**
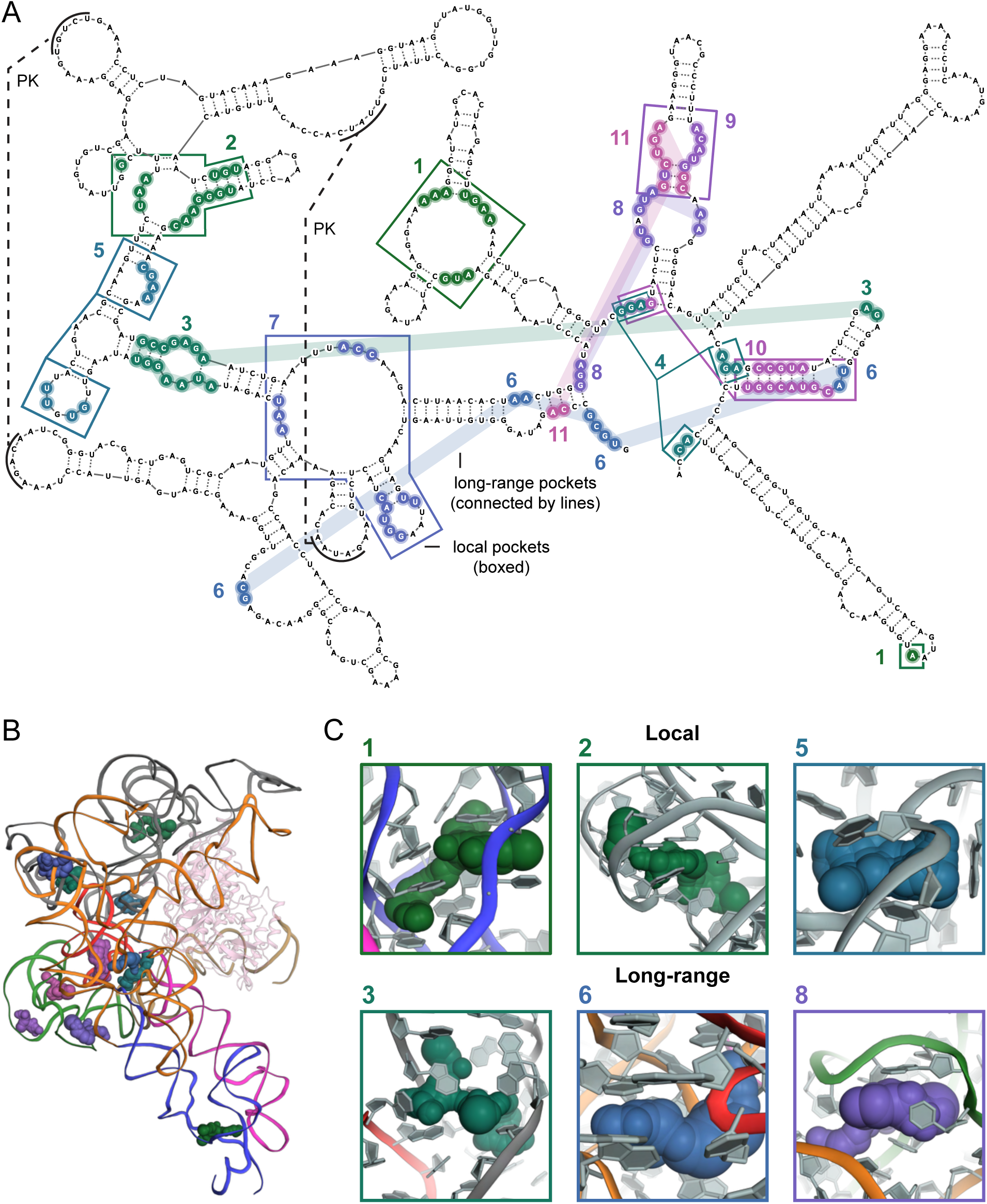
Representative illustration of pockets formed by local versus long-range interactions, detected in a group II intron RNA. Pockets formed by local domains are boxed. (*A*) Pockets identified by number and color, and superimposed on the RNA secondary structure. (*B*) Pockets visualized in the three-dimensional RNA structure (colored by domain). Complex contains a reverse transcriptase cofactor (pink) (PDB: 5G2X) (55). (*C*) Examples of pockets formed by local versus long-range RNA interactions. RNA backbone is colored by domain.

Pockets formed by long-range interactions in large RNA structures would be difficult or impossible to identify from a secondary structure model alone. Thus, we focused our analysis on local pockets. Small RNAs (< 200 nucleotides) form local pockets exclusively. A majority (66%) of pockets in multi-domain RNAs also occur in local secondary structures (**Fig. 7C**), which emphasizes that readily characterized secondary structure motifs form pockets in both small and large RNAs. We assigned each pocket detected by *fpocketR* to a class defined by the most complex secondary structural motif that contributes to pocket formation. Secondary structure motifs that formed pockets varied in complexity and included (*i*) simple loops and bulges, (*ii*) consecutive loops or bulges separated by a contact distance of 5 nucleotides or less, (*iii*) multi-helix junctions, and (*iv*) pseudoknots.

Simple bulges and loops are, as expected, extremely common and we observed 402 in our dataset; only 3% of these motifs formed a pocket. In addition, we observed that pockets in simple RNA motifs typically did not form independently but tended to form in large, multi-domain RNAs, where the global architecture could stabilize a local pocket in a simple structure (**Fig. 8A**). Pockets were slightly more common among consecutive loops (4% of which formed pockets; among 273 examples).

**Figure 8.**
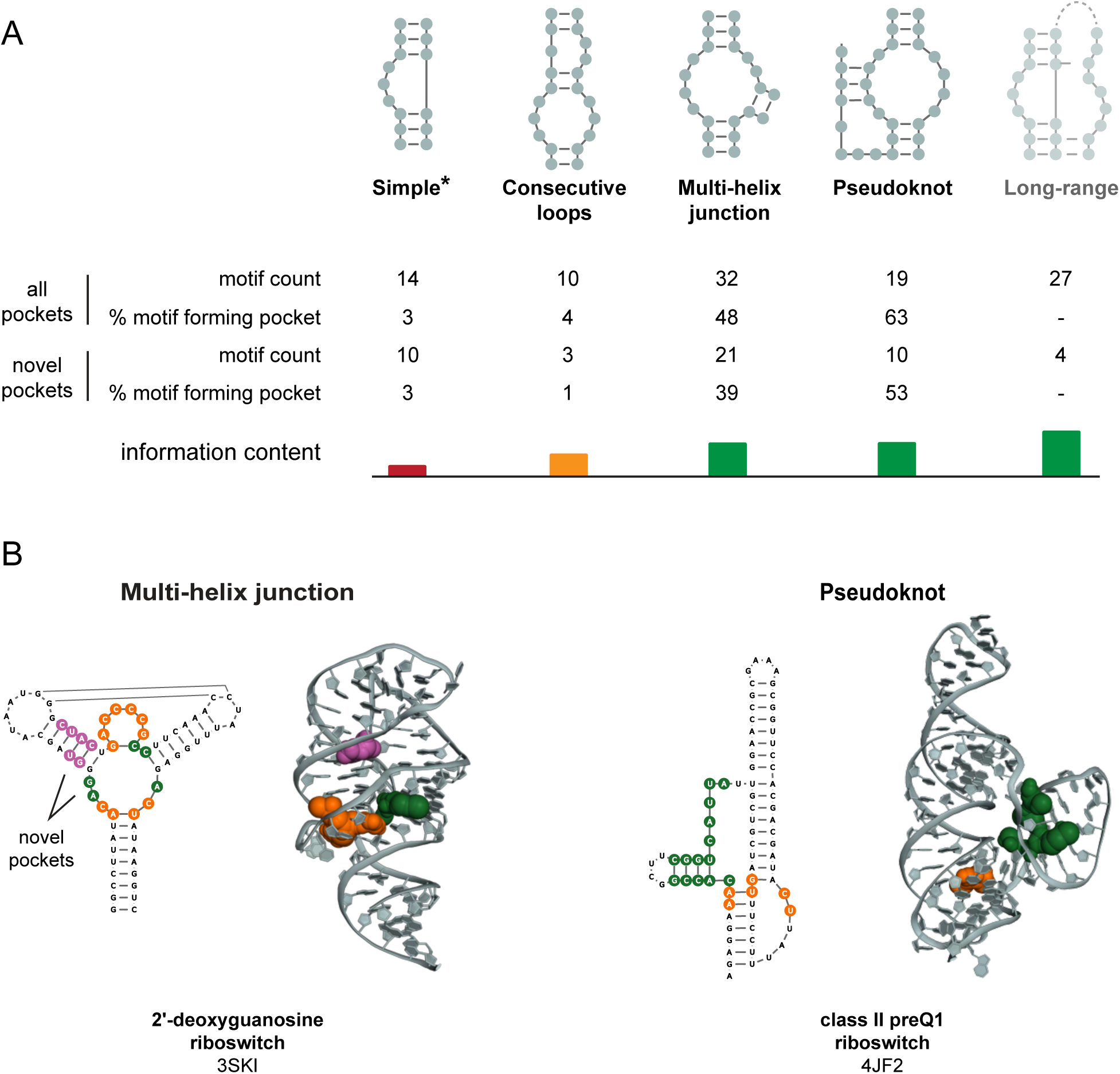
Secondary structural motifs that form pockets able to bind ligands with (drug-like) high-quality physicochemical properties. (*A*) Likelihood of secondary motifs to form pockets, in ascending order of complexity and information content. *, simple pockets were observed primarily in large multi-domain RNAs. (*B*) Examples of novel pockets in RNA secondary structural motifs most likely to form pockets. Known pockets are orange; novel pockets are green and magenta.

Pockets were much more common among complex structures, multi-helix junctions and pseudoknots, based on our all-pockets dataset (48% and 63% of which formed pockets, respectively) (**Fig. 8A**). Across our full dataset, the number of multi-helix junction and pseudoknot motifs is artificially increased, due to the inclusion of many riboswitch and aptamer motifs. Therefore, we additionally determined the local structure among only our novel pockets. Simple motifs and consecutive loops very rarely formed novel pockets (3% and 1%, respectively). Multi-helix junctions and pseudoknots formed novel pockets at high rates (39% and 53%, respectively) (**Fig. 8A**), comparable to those in the all-pockets dataset. This analysis emphasizes that any given simple motif is unlikely to contain a pocket able to bind small molecules with good physicochemical properties; but that many complex multi-helix junction and pseudoknot motifs will.

## DISCUSSION

A very small number of compounds, with plausibly drug-like physicochemical properties, have ever been visualized bound to compact RNA motifs at high resolution. A few additional drug-like (and actual drug) compounds have been visualized bound to the ribosome. The sizes of our training and test sets were therefore necessarily small, but the selection of these sets is a strength, not a weakness, of our approach. The field is not quite ready for machine learning to characterize these sites. *fpocketR* was optimized stringently using RNAs in complex with drug-like ligands (**Fig. 1**). We used the QED framework (21), which integrates eight physical features strongly associated with druglikeness into a single metric. We required ligands to have QED values greater than 0.3, which tends to eliminate non-selective RNA binders. Our analysis of antibiotic binding pockets in the ribosome found that *fpocketR* selectively detected pockets that bind drug-like ligands and, critically, distinguishes between pockets that bind drug-like versus un-drug-like ligands.

We were then able to leverage *fpocketR* to substantially expand the number of identified pockets able to bind drug-like ligands. In principle, RNA-targeted ligand discovery could open broad opportunities for manipulating gene expression, especially for targeting difficult-to-ligand proteins and non-coding RNAs. A critical starting point is to understand the rules that underlie RNA targets containing pockets able to bind small molecule ligands with good physicochemical features.

When applied to RNA, the protein-optimized *fpocket* algorithm (mis)identified numerous near- or non-pockets and, moreover, did not accurately rank accepted pockets. Ultimately, *fpocket*, with default parameters, did not perform well with RNA, emphasizing that fundamental differences exist between the ligand-facing surfaces of proteins and RNA. We identify three broad differences between pockets in RNA and protein. First, pockets detected in RNA (using *fpocketR*) and protein (using either algorithm) have broadly similar physical properties with the critical difference that pockets in RNA are more polar and less hydrophobic (**Table S10**).

Second, electrostatic properties significantly influence the pocket scoring functions used to rank pockets by their quality, and the distinctive electrostatics of RNA cause true pockets to be mis-ranked using current algorithms. Third, the ubiquitous presence of solvent-exposed major and minor grooves on the surface of RNA creates pseudo-pockets. False pocket exclusion by *fpocketR* was achieved by reducing reliance on pocket scoring functions to rank pockets and by imposing stricter geometric requirements that penalize highly solvent-exposed pockets.

We validated predictions made by *fpocketR* with Frag-MaP, which is unique relative to current approaches for detecting RNA-ligand interactions in complex mixtures (29, 35–37). Frag-MaP leverages mutational profiling (MaP) to identify RNA-ligand binding sites with high sensitivity and independently from biases introduced by biotin pulldown and adapter-ligation steps in seq-based experiments (24, 38). Here we used Frag-MaP in a simple way, to detect fragment binding sites in abundant rRNAs. Ligand-binding sites identified by Frag-MaP consistently surrounded pockets detected by *fpocketR*, demonstrating the general ligandability of novel pockets detected by *fpocketR.* A powerful future application will be to use Frag-MaP as an in-cell RNA tertiary structure discovery tool because ligand-binding pockets form preferentially and specifically in regions with complex RNA structures.

Analyses enabled by *fpocketR* emphasize that many pockets are created by nucleotides that are localized in RNA secondary structure. Local RNA secondary structures harbor nearly all the pockets in small RNAs (< 200 nucleotides) and form most pockets in large multi-domain RNAs. Pockets in secondary structures are primarily formed by complex motifs, including multi-helix junctions and pseudoknots, compared to simpler RNA structures such as bulges and loops.

Complex structural motifs both have a much higher rate of forming drug-like pockets and also have higher information content, which facilitates selective interactions with drug-like ligands across RNA transcriptomes (3) (**Fig. 8A**). For riboswitches, ligand binding sites were noted to form from residues far away in sequence but close in three-dimensional space, corresponding to regions of maximum contact order (39). Similarly, the multi-helix junctions and pseudoknots evaluated here have the highest rates of forming pockets and the largest contact orders.

Despite these clear advantages of targeting complex structures, a significant component of current work directed at discovery of RNA-targeting small molecules focuses on RNA motifs with simple structures (reviewed in (3, 18–20)). Our study indicates that simple bulge and consecutive loop motifs contain pockets only infrequently (**Fig. 8A**). These observations emphasize that RNA secondary structure is a useful tool to estimate RNA targetability and that RNAs with complex structural motifs are likely the best targets for drug-like small molecules (**Fig. 8B**).

*De novo* design of small-molecule ligands that target RNA and modulate function remains challenging. This work provides critical computational and experimental tools for identifying and understanding features of RNA structures that form targetable pockets, specifically able to bind drug-like small molecule ligands with high potential for therapeutic development.

## MATERIALS AND METHODS

### Curation of RNA-ligand complexes containing ligands with favorable (drug-like) physicochemical properties

RNA structures used to optimize *fpocketR* were obtained from the HARIBOSS non-redundant RNA-ligand database (11). RNA-ligand complex structures were required to (*i*) bind to a drug-like small molecule ligand (QED score ≥ 0.3), (*ii*) be high-resolution (≤ 3.5 Å), (*iii*) be a unique RNA in our data set, and (*iv*) ≤ 200 nucleotides (this requirement removes ligands that bind the ribosome). The quantitative estimate of drug-likeness (QED) score (21) was calculated using the QED module in the rdkit library (40). We emphasize that the QED metric is an open source and highly useful measure of fundamental physicochemical properties: molecular mass, hydrophobicity, hydrogen bond donors and acceptors, polarizable surface area, number of rotatable bonds, number of aromatic rings, and structural alerts (21). As defined by these criteria, there were 48 known RNA-ligand complexes with plausible drug-like ligands, that bind compact RNA sites. These RNA-ligand complexes were manually evaluated in PYMOL (41) to assess the solvent accessibility and cavity depth of the ligand binding pocket. Ultimately, 38 unique RNA structures were chosen and randomly split into a training set (n=13) and test set (n=25). For analysis of paired ligand-bound and apo structures (**Fig. 5A** and **Table S5**), we assessed all examples (10 total) of small RNA structures (< 500 nt) with published holo (bound to small-molecule ligands) and apo structures available in the PDB at the time of our study.

### Pocket evaluation metrics

Only pockets with an *fpocket* score ≥ 0 were analyzed. Pockets identified by *fpocket* were required to meet three criteria to be classified as a known pocket that binds a small-molecule ligand. The geometric center criterion is met if the geometric center of a pocket is within 4.5 Å of any atom of the ligand. The ligand overlap criterion is met if at least 25% of the alpha spheres in a pocket overlap the ligand (within 3 Å). The pocket overlap criterion is met if at least 25% of the atoms in the ligand overlap with a pocket (within 3 Å). Pockets that do not meet all three criteria are classified as novel pockets. However, we note that several novel pockets are occupied by biomolecules and ions (**Table S6-S9**). Software used to evaluate these criteria are provided (see data and software availability section).

### Multivariate optimization of *fpocket*

We optimized *fpocket* 4.0.3 (22) for detecting pockets in RNA by systematically adjusting values of six core parameters (*M*, *m*, *D*, *i*, *A*, *p*). Parameters A and p filter pockets based on polarity and, in preliminary analyses, were not effective at improving pocket detection in highly polar RNA molecules. The remaining four parameters (*m*, *M*, *D, i*) control the minimum (*m*) and maximum (*M*) alpha sphere radii, clustering distance (*D*), and minimum cluster size (*i*) (**Fig. 1A**) and were systematically optimized in 3 rounds of multivariate optimization using our training set of RNA-ligand complexes (*n* = 13). We evaluated *fpocket* performance to detect ligand binding sites in a large parameter space (1820 parameter combinations) by performing a coarse, medium, and fine round of optimization (**Table S3**). In each round, we evaluated 5 values for each parameter with progressively smaller increments between the values. We started with the default *fpocket* parameters and their flanking values and used the best performing parameters from each round of optimization to generate values for the subsequent round. The best performing parameters were selected by calculating a pareto set that maximized sensitivity and positive predictive value (ppv) and by applying some intuition intended to avoid overtraining (42). We also propose alternative relaxed parameters for investigation of pockets and near-pockets that lie just outside of these criteria (see Supporting Information).

### fpocketR usage

RNA pockets were detected using *fpocketR*. *fpocketR* is a wrapper for *fpocke*t 4.0.3 (22) that provides options for advanced analysis and visualization of pockets detected in RNA structures. RNA pockets were detected using the input --pdb option for a .pdb or .cif file, the --chain option (maximum of 2 RNA chains), and the newly defined *fpocketR* parameters (--*m* 3.0, --*M* 5.7, --*D* 1.65, --*i* 42). Multistate NMR ensembles were analyzed using the --state 0 and - -qualityfilter 0.4 options. Known binding ligands were specified using the --ligand and -- ligandchain options, allowing *fpocketR* to differentiate known pockets (visualized in output files as orange to yellow) from novel pockets (green to pink). Visualizations for pockets in secondary structure space were generated by inputting a secondary structure drawing using the --nsd and --connectpocket options.

### Pocket visualization

Figures visualizing predicted pockets were generated using the *fpocketR* software package. *fpocketR* generates tertiary structure figures using the *PyMOL* API (41) and secondary structure figures using *RNAvigate* (43). *fpocketR* visualizes pockets as a group of alpha-atoms (computed by subtracting 1.65 Å from the *fpocketR*-generated alpha sphere radius) (Fig. 1A). Pockets are visualized in secondary structures by coloring the nucleotides that contact alpha spheres from a pocket. RNA secondary structure drawing templates (.nsd) were generated from PDB structures using RNApdbee (44) manually edited in StructureEditor (45), and input into *fpocketR* using the --nsd option.

### Classifying pocket structural motifs

Pocket-forming secondary structures were classified by the types of RNA structure in contact with alpha spheres. The classes identified and used in this work were, in order of structural complexity: simple structures, consecutive loops, multi-helix junctions, pseudoknots, and long-range structures. Specific structural definitions were: Simple structure, pocket only contacts nucleotides in an apical loop, internal loop, or bulge. Consecutive loops, pocket contacts a pair of loops/bulges within a contact distance of 5 nucleotides from each other (contact distance is defined as the shortest path length through the secondary structure graph between two nucleotides). Multi-helix junction, pocket contacts single-stranded nucleotides in the junction or base-paired nucleotides within 3 nucleotides of the multi-helix junction. Pseudoknot, pocket contacts base-paired nucleotides involved in a non-nested interaction or single-stranded nucleotide located between non-nested helices. Long-range structure, pocket contacts motifs in a structure that have a contact distance of >15 nucleotides. If a structure fit the criteria for multiple motif classifications, the structure was assigned to the most complex structural motif. Structures (4 observed) containing g-quadruplexes were not analyzed.

### In-cell Frag-MaP

Experiments were performed in *Bacillus subtilis* subsp. *subtilis* strain 168 (ATCC) in LB at 30 °C. Cell were probed with 200 µM fragment probe or control probe for 30 min in the dark, placed on ice, and then exposed to 3 J/cm2 of 365-nm-wavelength UV light for 9 minutes (Analytik Jena). RNA was extracted and subjected to two bead-based clean up steps (silica-based SPE, Monarch RNA Cleanup Kit; NEB; and click chemistry-based, Magnefy Azide, medium 1.0 µm beads; Bangs Laboratories). RNA was then subjected to mutational profiling reverse transcription (MaP-RT) with random nonamer primers to generate cDNAs (46). Extended and detailed methods are provided in the Supporting Information.

### Sequencing and Data Analysis

cDNAs were sequenced (NEBNext Ultra II DNA Library Prep Kit for Illumina, NEB) using standard methods (46). Ligand-binding sites were identified using a new software package, *FragMapper*, implemented in *RNAvigate* (v1.0) (43). Frag-MaP sites were visualized in RNA tertiary structures using *PyMOL* (41). These steps are described in detail in the Supporting Methods.

## DATA AVAILABILITY

All data and software generated or used in this work are freely available. The *fpocketR* datasets reported in this study and *fpocketR* software suite and user manual are available at https://github.com/Weeks-UNC/2025_Veenbaas and https://github.com/Weeks-UNC/fpocketR, respectively. *FragMapper* software is available as an analysis module in RNAvigate, https://github.com/Weeks-UNC/RNAvigate. Frag-MaP datasets have been deposited in the Gene Expression Omnibus (GEO) database, https://www.ncbi.nlm.nih.gov/geo (accession no. GSE276279).

## ACKNOWLEDGEMENTS

This work was supported by NIH grant R35 GM122532 (to K.M.W.); software development for tertiary structure discovery by Frag-MaP was supported by the NSF (MCB-2027701 to K.M.W.). J.T.K. is a UNC Lineberger Integrated Training in Cancer Model Systems Fellow (T32 CA009156) and an NIH Kirschstein Postdoctoral Fellow (F32 GM143863). We are indebted to Rachel M. Johnson (University of North Carolina) for performing high-resolution mass spectrometry experiments. We are grateful to members of the Weeks Laboratory, especially to Drs. Simon Felder and Alexandra Khitun, for their continued mentorship and thoughtful input to this work.

