## Supporting Information for "Ligand-binding pockets in RNA, and where to find them"

Seth D. Veenbaas, Jordan T. Koehn, Patrick S. Irving, Nicole N. Lama and Kevin M. Weeks*
Department of Chemistry, University of North Carolina, Chapel Hill NC 27599-3290

Supporting Materials and Methods, Supporting Figures 1-7, Supporting Tables 1-11, and Supporting References (including all examined RNA and RNA-ligand structures).

### Materials and Methods

**General chemistry information.** Reactions were carried out in amber glass sample vials. All reagents, starting materials, and solvents (including dry solvents) were obtained from commercial suppliers and used without further purification. Thin-layer chromatography (TLC) was performed using commercial silica gel 60 $F_{254}$ coated aluminum-backed sheets; products were visualized with UV light. Purification was carried out by automated flash chromatography (Selekt; Biotage) using normal phase columns (Sfär; Biotage). All NMR ($^1$H and $^{13}$C) spectra were recorded at 400 MHz with a dual carbon/proton cryoprobe, or at 600 MHz with a dual carbon/proton cryoprobe. NMR samples were recorded in $CDCl_3$ or $CD_3OD$. Chemical shifts are reported in parts per million (ppm) and referenced to the center line of residual solvent (for $CDCl_3$, $\delta$ 7.26 ppm for $^1$H NMR and 77.16 for $^{13}$C NMR; for $CD_3OD$, $\delta$ 3.31 for $^1$H NMR and 49.0 ppm for $^{13}$C NMR). Coupling constants are reported in Hertz (Hz). High-resolution mass spectrometry was performed on an Agilent 1200 series analytical high performance liquid chromatography system coupled to an Agilent 6520 Accurate Mass quadrupole time of flight (Q-TOF) mass spectrometer and an electrospray ion source (ESI). Fragment 206 was obtained from Enamine.

**General synthesis procedure for fully functionalized fragment probes.** All compounds were synthesized following a general amide coupling procedure (1). 3-(3-(but-3-yn-1-yl)-3H-diazirin-3-yl)propanoic acid (1 eq, 60 mM), DIPEA (3.0 eq), EDC-HCl (1.5 eq) and HOBt (1.5 eq) were dissolved in DCM and added to a commercially available amine (1.1 eq). Reaction mixtures were stirred at 20 ºC overnight and monitored by TLC. The crude product was diluted in DCM (10 mL), washed with saturated aqueous $NH_4Cl$ (10 mL), and then washed with saturated aqueous $NaHCO_3$ (10 mL). The organic layer was dried over anhydrous $Na_2SO_4$ and solvent removed by rotary evaporation under reduced pressure. The crude products were purified by PTLC or flash column chromatography (Biotage).

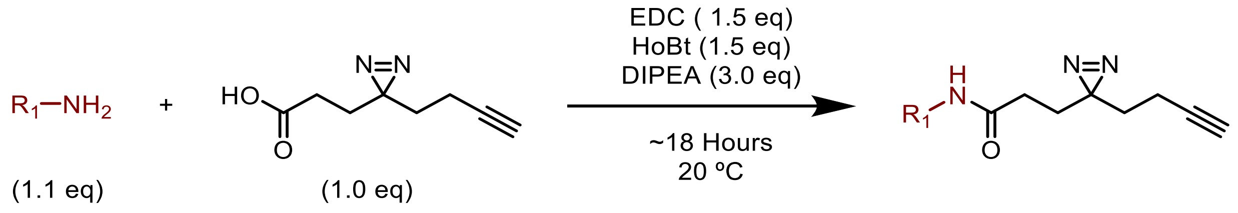

**Linezolid probe: (R)-3-(3-(but-3-yn-1-yl)-3H-diazirin-3-yl)-N-((3-(3-fluoro-4-morpholinophenyl)-2-oxooxazolidin-5-yl)methyl)propenamide.** Prepared using the general synthesis procedure and (R)-5-(aminomethyl)-3-(3-fluoro-4-morpholinophenyl)oxazolidin-2-one (39 mg, 0.13 mmol). Purified by $SiO_2$ flash chromatography on a Biotage (DCM/MeOH, 100:0 to 99:1) yielding 13.2 mg of a white solid (25%). $^1$H NMR (600 MHz, Methanol-$d_4$) δ: 7.51 (dd, $J$ = 14.7, 2.6 Hz, 1H), 7.18 (ddd, $J$ = 8.8, 2.6, 1.1 Hz, 1H), 7.07 (t, $J$ = 9.2 Hz, 1H), 4.80 (dddd, $J$ = 9.0, 6.2, 5.3, 4.2 Hz, 1H), 4.13 (t, $J$ = 9.0 Hz, 1H), 3.85 (t, $J$ = 6.3 Hz, 1H), 3.84 (t, $J$ = 4.4 Hz, 4H), 3.62 (dd, $J$ = 14.5, 5.2 Hz, 1H), 3.51 (dd, $J$ = 14.5, 4.0 Hz, 1H), 3.06 (t, $J$ = 4.4 Hz, 4H), 2.26 (t, $J$ = 2.7 Hz, 1H), 2.07 – 2.00 (m, 2H), 1.95 (td, $J$ = 7.5, 2.7 Hz, 2H), 1.71 (td, $J$ = 7.5, 3.0 Hz, 2H), 1.58 – 1.52 (m, 2H). $^{13}$C NMR (151 MHz, MeOD) δ: 137.3, 135.2, 135.1, 130.2, 126.7, 115.4, 110.7, 108.5, 108.3, 86.9, 83.6, 73.5, 70.4, 67.9, 52.4, 52.4, 49.4, 49.3, 49.1, 49.0, 48.9, 48.7, 48.6, 42.9, 33.3, 30.9, 29.7, 28.8, 13.8. HRMS (ESI-QTOF) calculated $[M+H]^+$ = 444.2042; observed m/z = 444.2057.

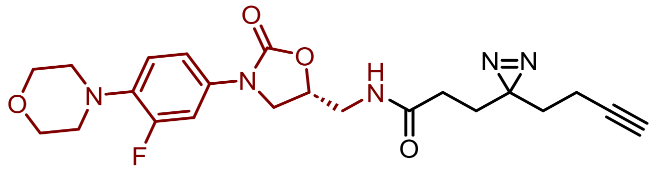


**Methyl probe: 3-(3-(but-3-yn-1-yl)-3H-diazirin-3-yl)-N-methylpropanamide.** Prepared using the general synthesis procedure and methylamine HCl (8.9 mg, 0.13 mmol). Purified by $SiO_2$ flash chromatography on a Biotage (Hexanes/EtOAc, 7:3 to 1:1) yielding 15.0 mg of colorless solid (70%). $^1$H NMR (400 MHz, Chloroform-$d$) δ: 5.51 (s, 1H), 2.81 (d, $J$ = 3.9 Hz, 3H), 2.05 – 1.97 (m, 3H), 1.92 (ddd, $J$ = 8.0, 6.2, 1.8 Hz, 2H), 1.85 (ddd, $J$ = 8.7, 6.3, 1.8 Hz, 2H), 1.65 (t, $J$ = 7.4 Hz, 2H). HRMS (ESI-QTOF) calculated $[M+H]^+$ = 180.1131; observed m/z = 180.1130.

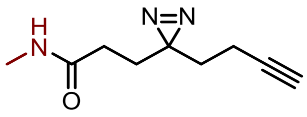


**6-Aminoquinoxaline probe: 3-(3-(but-3-yn-1-yl)-3H-diazirin-3-yl)-N-(quinoxalin-6-yl)propenamide.** Prepared using the general synthesis procedure and quinoxaline-6-amine (19 mg, 0.13 mmol). Purified by PTLC (Pentane/EtOAc, 1:4) yielding 4.5 mg of a pale-yellow oil (13%). $^1$H NMR (400 MHz, Methanol-$d_4$) δ 8.79 (dd, J = 19.2, 1.9 Hz, 2H), 8.52 (d, J = 2.3 Hz, 1H), 8.02 (d, J = 9.1 Hz, 1H), 7.92 (dd, J = 9.1, 2.4 Hz, 1H), 2.34 – 2.24 (m, 3H), 2.06 (td, J =

7.5, 2.7 Hz, 2H), 1.89 (dd, J = 8.4, 6.9 Hz, 2H), 1.66 (t, J = 7.4 Hz, 2H). $^{13}C$ NMR (101 MHz, Methanol-$d_4$) δ 173.16, 146.77, 144.98, 144.63, 141.85, 141.00, 130.43, 125.49, 117.34, 83.60, 70.35, 33.50, 31.93, 29.38, 28.89, 13.86. HRMS (ESI-QTOF) calculated $[M+H]^+$ = 294.1349; observed m/z = 294.1348.

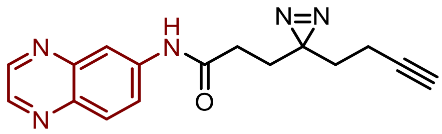


**Comparison of protein and RNA pocket detection and properties.** Protein structures (n = 200) were randomly selected from a curated list of protein-ligand complexes (2) and acquired from the PDB (3). All RNA structures in the training and test sets of this work were used. Pocket prediction and characterization for both protein-ligand and RNA-ligand complexes was performed using *fpocketR* with default (or *fpocket*) parameters defined as ($m$ = 3.4, $M$ = 6.2, $D$ = 2.4, and $i$ = 15) and optimized (or *fpocketR*) parameters defined as ($m$ = 3.0, $M$ = 5.7, $D$ = 1.65, and $i$ = 42).

**Alternative, relaxed, parameters for *fpocketR*.** In some instances, it is useful to employ a "relaxed" parameter set to explore additional possible pockets and near-pockets in an RNA. Figure S1 illustrates a region of strongly-performing values at the intersection of high sensitivity and high PPV corresponding to the parameter ranges: $m$: 2.9 – 3.1, $M$: 5.5 – 5.8, $D$: 1.5 – 1.7, and $i$: 36 – 44, and. For useful relaxed parameters, we specifically recommend decreasing the number of alpha-spheres required for a pocket (--i 35) and increasing the maximum alpha size (--M 5.8).

**In-cell probing for Frag-MaP.** *Bacillus subtilis* subsp. *subtilis* strain 168 (ATCC) was grown on LB agar plates at 30 ºC. A 5 mL culture was inoculated with a single colony and grown overnight in LB media at 30 ºC, 225 rpm. A 2 mL aliquot of the culture was diluted (1:25) with LB media to create a subculture that was grown at 30 ºC, 225 rpm to $OD_{600}$ ~0.5. The cells were pelleted at 3,000 x*g* for 10 minutes, washed once with PBS, pelleted again, and then resuspended in a final volume of 25 mL PBS. Resuspended cells (5 mL) were then incubated with 200 µM fragment probe or control probe for 30 minutes in the dark at 30 ºC, 225 rpm. Treated cells (5 mL) were then transferred to 6-well plates, placed on ice, and exposed to 3 J/cm2 of 365-nm-wavelength UV light over 9 minutes (Analytik Jena UVP CL-1000 equipped with five 8-W F8T5 black lights) at 10 cm from the light source. Cells were collected and pelleted at 3,000 x*g* for 10 minutes, resuspended in 250 µL lysis buffer (30 mM Tris pH 7.0, 10 mM EDTA, 10 mg/mL lysozyme), and incubated at 25 ºC for 30 minutes. Total RNA was then extracted (1 mL of TRIzol; Invitrogen) and purified with on-column DNase treatment (Monarch Total RNA Miniprep; NEB).

**Click labeling and RNA enrichment.** Total RNA was chemically fragmented at 94 ºC for 2 minutes to yield RNAs with an average size of ~200 nucleotides (Magnesium RNA Fragmentation Module; NEB). RNA fragments were purified (silica-based SPE, using Monarch RNA Cleanup Kit; NEB). The RNA fragments (0.5 µM) were heat-denatured at 90 ºC for 3 minutes and "clicked" on to azide-linked beads (0.5 µm medium magnify beads; Bangs Laboratories, Inc.) with 240 µM copper(II)-TBTA complex (Lumiprobe), 500 µM fresh ascorbic acid (Sigma), and 50% v/v anhydrous DMSO (Invitrogen), in a final reaction volume of 50 µL. Click reactions were sparged with nitrogen, incubated at 40 °C for 30 minutes, and quenched with 1 µL 0.5 M EDTA. Cross-linked RNA was enriched by resuspending the RNA four times in wash buffer (1M NaCl, 10 mM Tris pH 7.5, 1 mM EDTA, 0.01% TWEEN-20) and a final time in nuclease-free water, the supernatant from each wash was removed via pipette after immobilizing the bead-conjugated RNA using a magnetic separation rack (NEB). Note that this process is used only for enrichment, not to call ligand-engagement sites.

**MaP reverse transcription.** Frag-MaP directly detects ligand crosslinking at RNA binding sites using mutational profiling, differentiating it from pull-down seq-based strategies. RNA was subjected to mutational profiling reverse transcription with random nonamer primers for rRNA analysis, a 5 min 90 ˚C denaturation step was added (4). Bead-conjugated RNA was mixed with 100 ng of random nonamer primer (NEB), and 20 nmol of dNTPs (NEB), and denatured at 90 ˚C for 5 minutes followed by incubation at 4 ˚C for 2 minutes. MaP-RT buffer (6 mM $MnCl_2$, 1 M betaine, 50 mM Tris (pH 8.0 at 25 ˚C), 75 mM KCl, and 10 mM fresh DTT) was added to the RNA solution and incubated at 25 ˚C for 2 minutes. SuperScript II Reverse Transcriptase (1 µL, 200 units; Invitrogen) was added and the reverse transcription reaction was performed according to the following temperature program: 25 °C for 10 min, 42 °C for 90 min, 10 × [50 °C for 2 min, 42 °C for 2 min], 72 °C for 10 min (4). The resulting cDNA was purified via magnetic pull-down using a magnetic separation rack (NEB). Frag-MaP can also be readily performed with gene-specific primers.

**Library preparation and sequencing.** Double-stranded DNA was prepared from the reverse transcription reaction product (NEBNext Ultra II Non-directional RNA Second Strand Synthesis Module; NEB) and DNA sequencing libraries were prepared (NEBNext Ultra II DNA Library Prep with Sample Purification; NEB). Libraries were then quantified (Qubit high-sensitivity dsDNA assay; Invitrogen and High sensitivity D1000 screentape; Agilent) and pooled as an equimolar sequencing library. The library was sequenced using 2x110 paired-end sequencing on an Illumina NextSeq 1000 instrument (P2 200 cycle, v3 chemistry; Illumina).

**Frag-Map Data analysis.** *ShapeMapper* (v2.1.5) was used to align reads to the 23S rRNA and calculate per nucleotide reactivities using the --random-primer-len option. The location of Frag-MaP sites was determined using the *FragMapper* analysis module written for *RNAvigate* (v1.0) (5). *FragMapper* can be used to quantify experiments executed in both random-primed and gene-specific modes (6). These modes enable Frag-MaP analysis (*i*) of RNA-ligand interactions transcriptome-wide without prior knowledge of the RNA target or (*ii*) of targeted gene regions to yield very high-quality confirmation of hits from transcriptome-wide screens. *FragMapper* requires *ShapeMapper* profiles for an RNA treated with a fragment probe and a fragment-less control probe. First, *FragMapper* filters out nucleotides that do not meet a minimum read depth threshold or are hyperreactive to the fragment-less (non-selective control) probe (mutation rate > 2.5%). Second, a modified z-score is calculated for each nucleotide based on the difference (delta) in mutation rates between the fragment-treated and methyl-treated (fragment-less) samples. Nucleotides with a significantly higher mutation rate for the fragment-treated sample (modified z-score > 30, z-score - standard error > 5, and delta mutation rate > 1%) are identified as Frag-MaP sites. Our analysis focused on RNA-only pockets; Frag-MaP sites detected near proteins in our reference structure (PDB: 7AS8) were not analyzed (**Fig. S4i**). Three Frag-MaP sites (2145, 2154, and 2198) were identified in the L1 stalk which is a highly conserved component of the ribosome but not resolved in crystal structures for the *B. subtilis* LSU. However, we found that the Frag-MaP sites in the L1 stalk were indeed close to pockets by analyzing a trRosettaRNA (7) model for the *B. subtilis* L1 stalk (eRMSD: 1.7 Å) (8) and a crystal structure of the L1 Stalk from Haloarcula marismortui (PDB: 5ml7) (9). Frag-MaP sites were visualized in RNA tertiary structures using *PyMOL* (10).

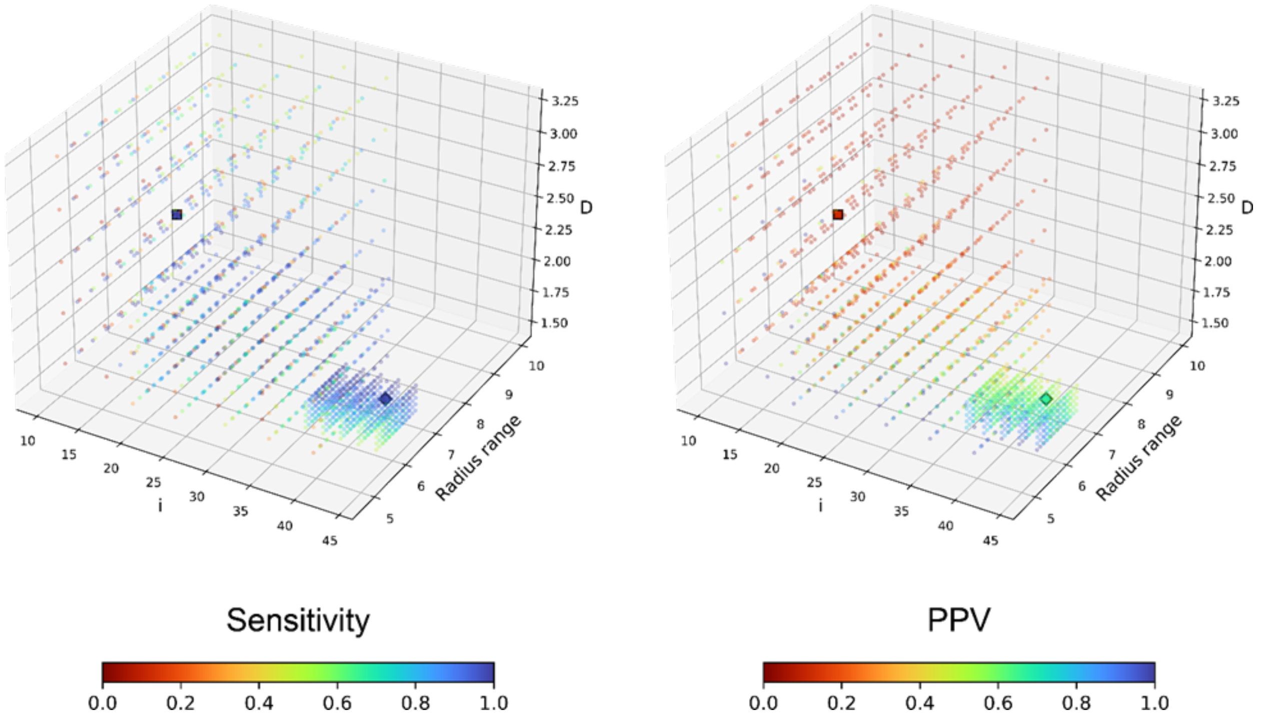


**Figure S1.** Parameter space evaluated during *fpocketR* optimization. Each point represents a unique combination of parameters (*i*, *m*, *M*, and *D*). Parameters *m* and *M* which control alpha sphere radius were collapsed to a single value representing radius range ($M^{1.3} – m$) to allow visualization in three-dimensions. Points are colored to reflect the sensitivity and positive predictive value (ppv) for detecting known ligand binding sites in the training set (n=13). The default (*i*=15, *m*=3.4, *M*=6.2, *D*=2.4) and optimized (*i*=42, *m*=3.0, *M*=5.7, *D*=1.65) parameters are depicted as squares and diamonds respectively.

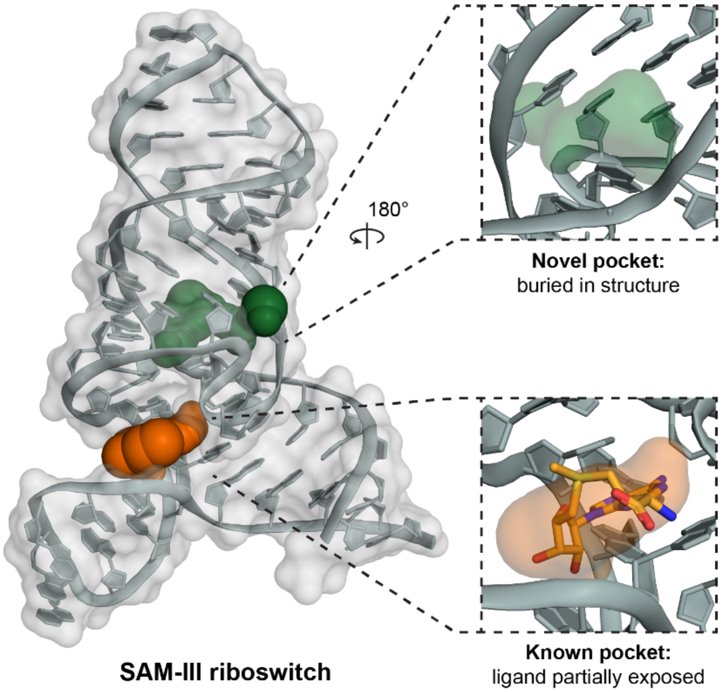


**Figure S2.** Pockets detected in the SAM-III riboswitch (PDB: 3E5C) (11) using *fpocketR*. The known pocket is formed in a three-way multi-helix junction and overlaps with most of the native SAM ligand but leaves the methionine tail exposed. The novel pocket forms between a three-way multi-helix junction and a bulge. Although the novel pocket is smaller in volume than the known pocket, it is more deeply buried in the RNA structure and has less solvent exposed surface area.

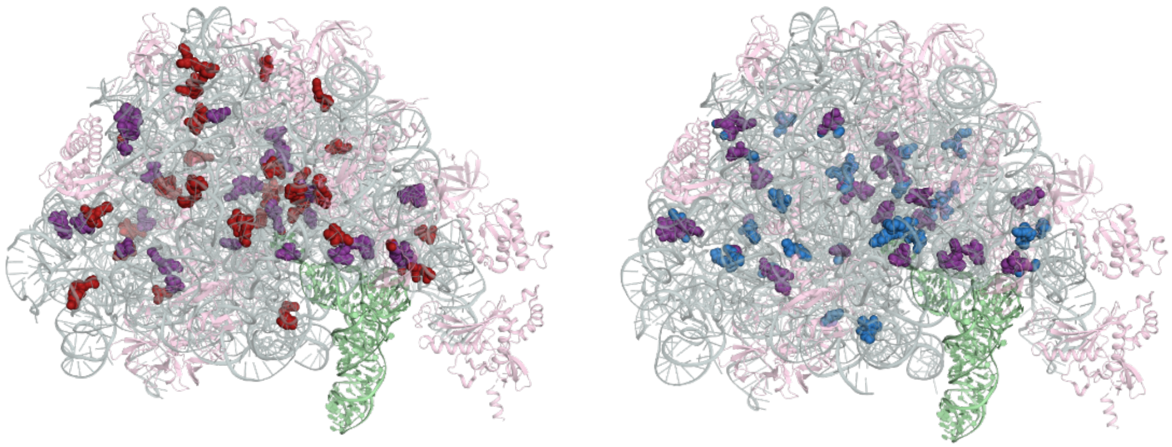

***E. coli* LSU**
(PDB: 7K00)

***B. subtilis* LSU**
(PDB: 7AS8)

**Figure S3.** The ribosomal pocket-ome. *fpocketR* detects 46 and 52 pockets in the classical (non-rotated) state of the 23S rRNA for *E. coli* and *B. subtilis* (12, 13), respectively. Of these, 21 pockets overlap between the two species (in purple). Ribosomal proteins are pink; A-site and P-site tRNAs are green.

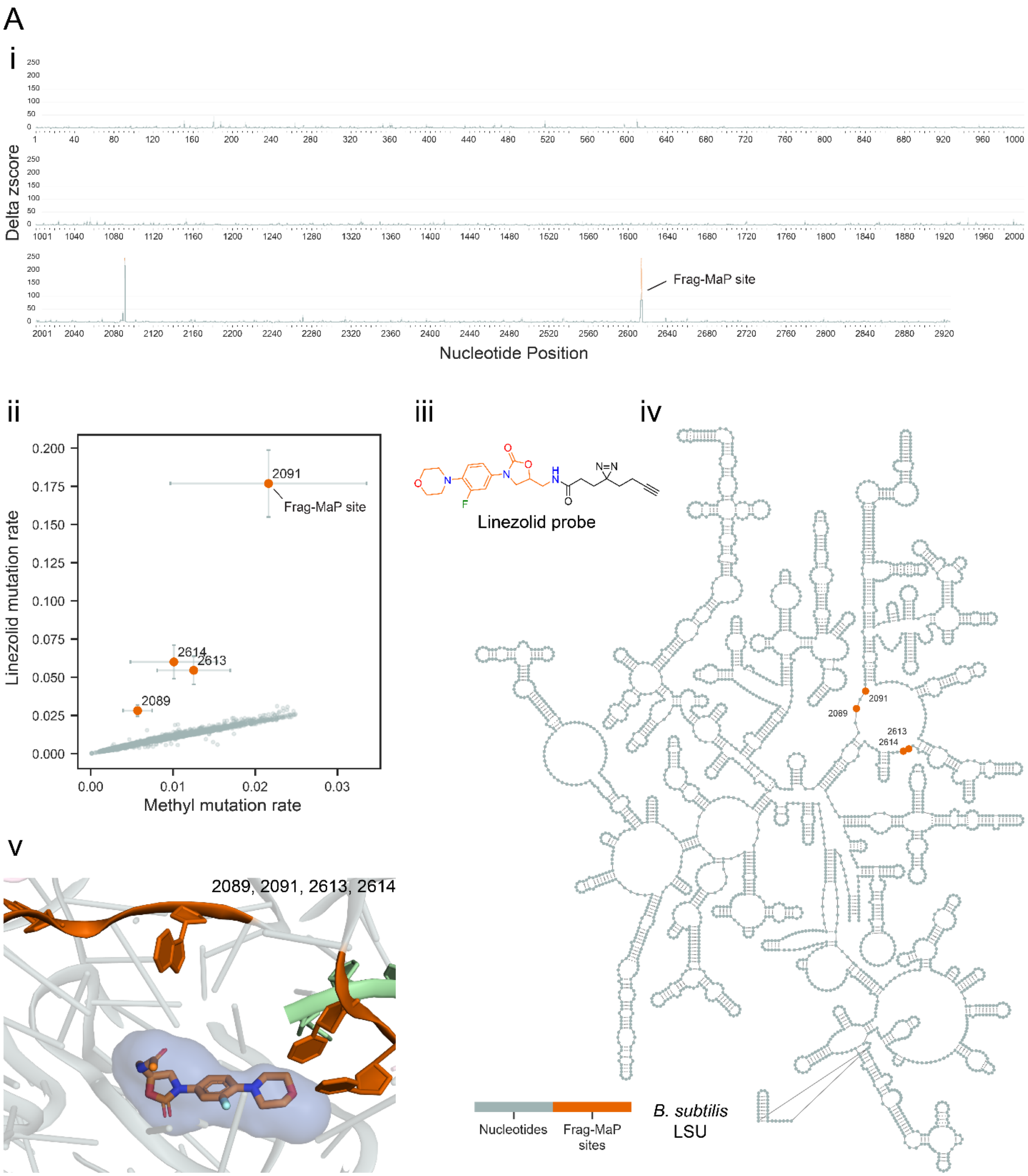
A
i
Delta zscore
Frag-MaP site
Nucleotide Position
ii
Linezolid mutation rate
Methyl mutation rate
2091
Frag-MaP site
2614
2613
2089
iii
Linezolid probe
iv
2091
2089
2613
2614
Nucleotides
Frag-MaP sites
B. subtilis LSU
v
2089, 2091, 2613, 2614

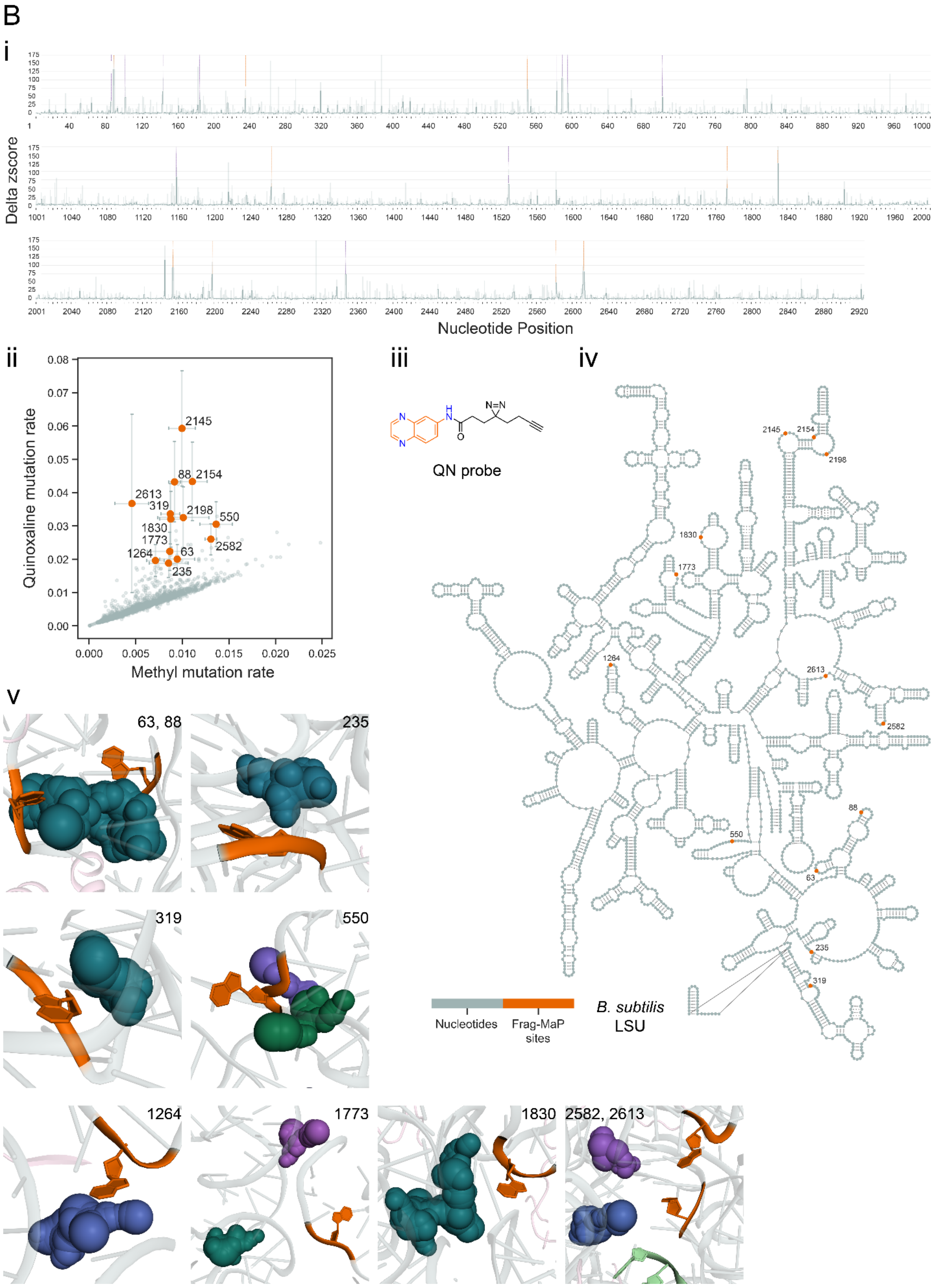
B
i
Delta zscore
Nucleotide Position
ii
Quinoxaline mutation rate
Methyl mutation rate
2145
88
2154
2613
319
2198
550
1830
1773
2582
1264
63
235
iii
QN probe
iv
B. subtilis
LSU
Nucleotides
Frag-MaP sites
v
63, 88
235
319
550
1264
1773
1830
2582, 2613

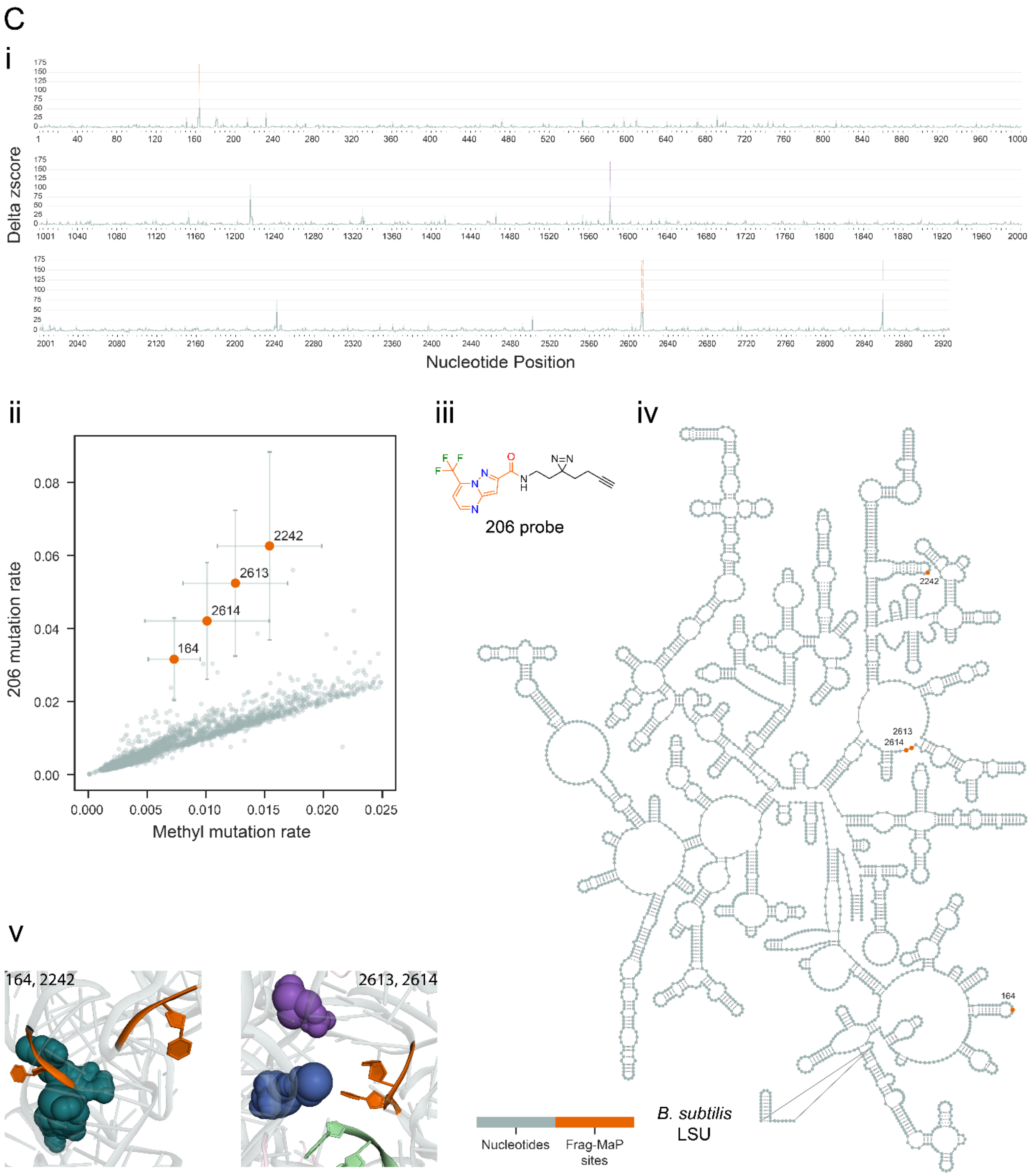
C
i
Delta zscore
Nucleotide Position
ii
206 mutation rate
Methyl mutation rate
2242
2613
2614
164
iii
206 probe
iv
2242
2613
2614
164
Nucleotides
Frag-MaP sites
*B. subtilis*
LSU
v
164, 2242
2613, 2614

**Figure S4.** (3 panels, *above*) Frag-MaP sites identified in the *B. subtilis* 23S rRNA using the (*A*) linezolid, (*B*) quinoxaline, and (*C*) 206 functionalized probes. For each panel: (*i*) Frag-MaP profiles displaying z-score of the difference in mutation rate between the fragment and methyl probes for each nucleotide. Frag-MaP sites analyzed in this work are depicted in orange. Frag-MaP sites at RNA-protein interactions or in structurally unvisualized regions, not analyzed in this work, are purple. Error is shown as the standard error of the mean. (*ii*) Comparison of the per-nucleotide mutation rate between the fragment and methyl probes. (*iii*) Structure of the functionalized fragment probe. (*iv*) Location of Frag-MaP sites (orange) within the secondary structure of the *B. subtilis* large subunit (LSU). (*v*) Proximity of Frag-MaP sites (orange) to pockets predicted by fpocketR (colored spheres) (PDB: 7AS8) (13).

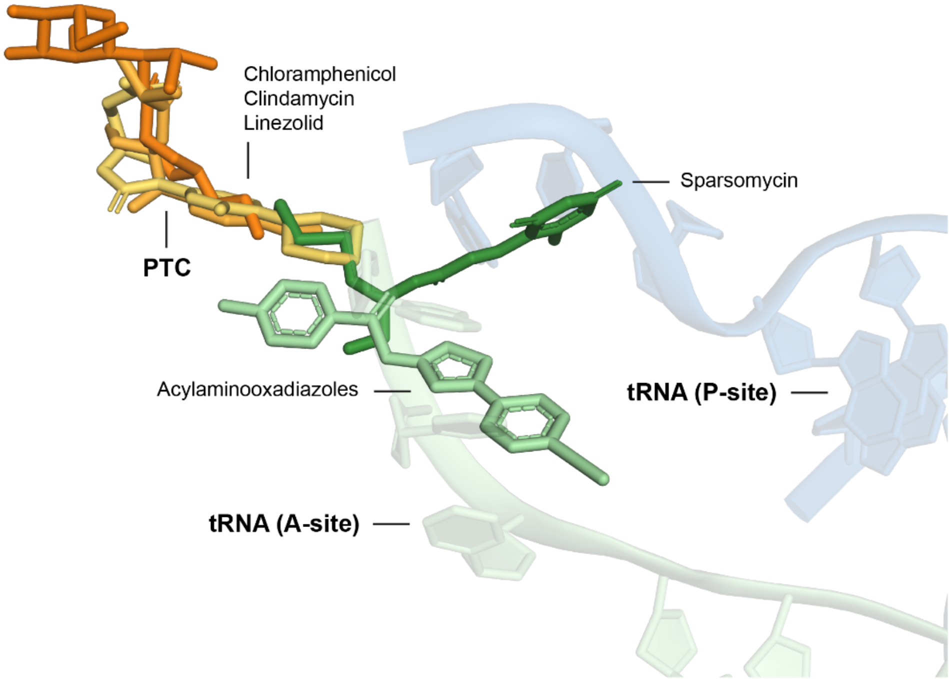


**Figure S5.** Distinct binding regions of ribosomal antibiotics near the peptidyl transferase center in the bacterial 23S rRNA. The binding position of chloramphenicol, clindamycin, and linezolid superimpose in the peptidyl transferase center. Sparsomycin and the acylaminooxadiazoles bind in an adjacent region overlapping the A-site tRNA. PDB codes are 6ND5, 4V7V, 3CPW, 1M90, and 6OM6 (14–18)

### Simple structures

1lvj_state3

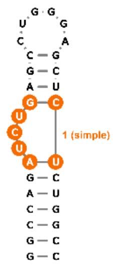


2ktz

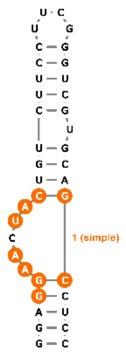


6va4

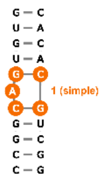


7fj0

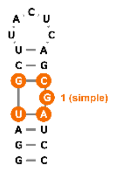


### Consecutive Loops

1q8n

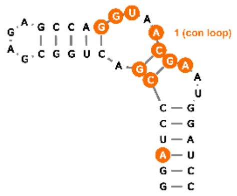


6xb7_state3

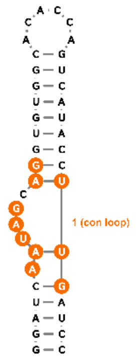


6xjq

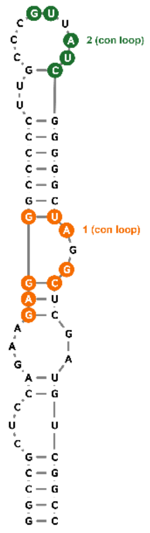


7dwh

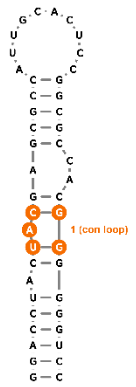


7elr

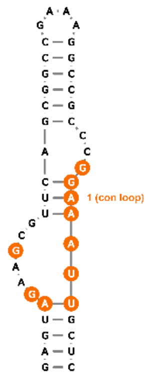


8d2b

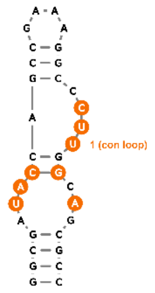

### Multi-helix Junctions

2gdi

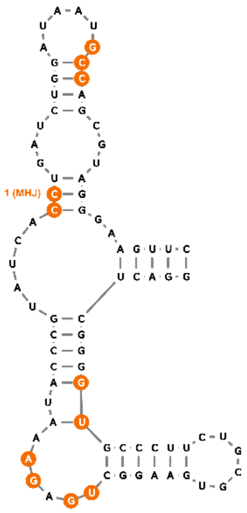


3e5c

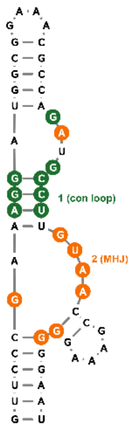


3d0u

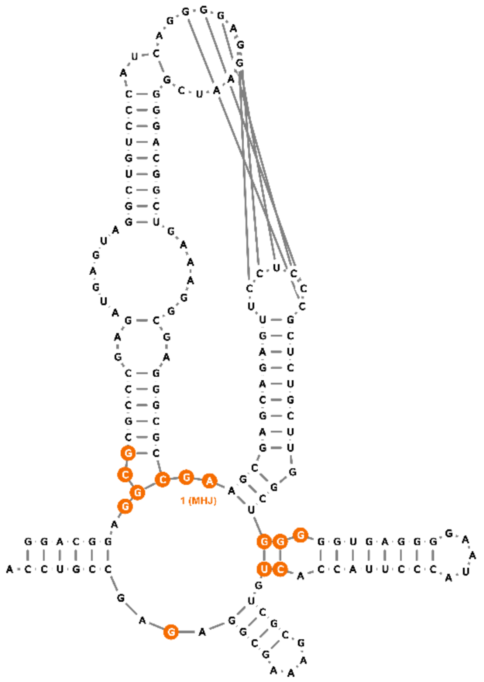


5kx9

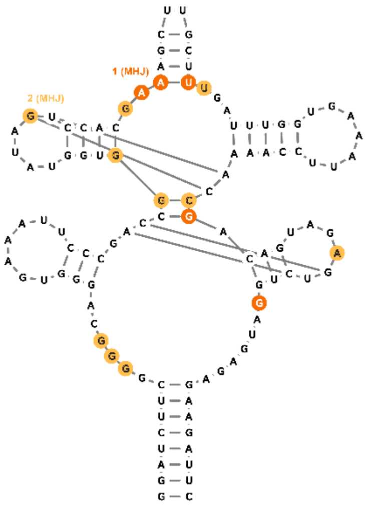


3ski

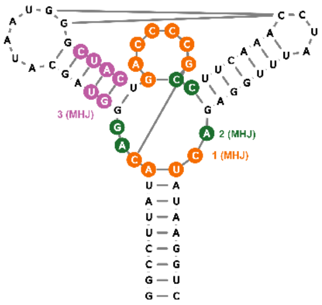


4b5r

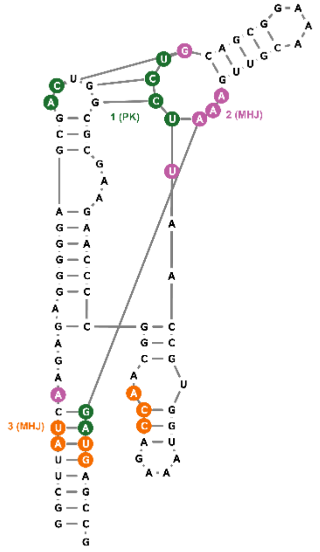

4lx5

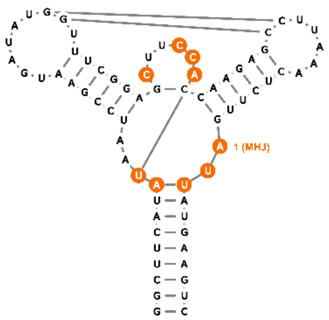


5kpy

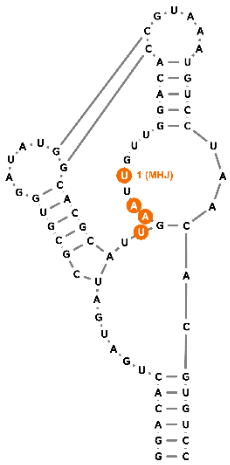


6gzr_state4

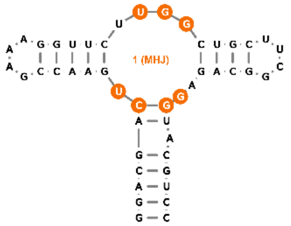


6las

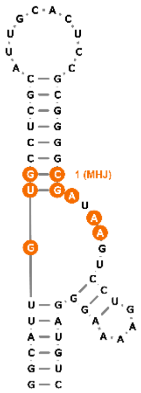


6ubu

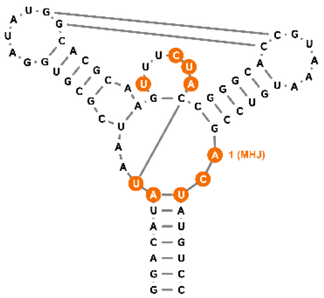


7kvt

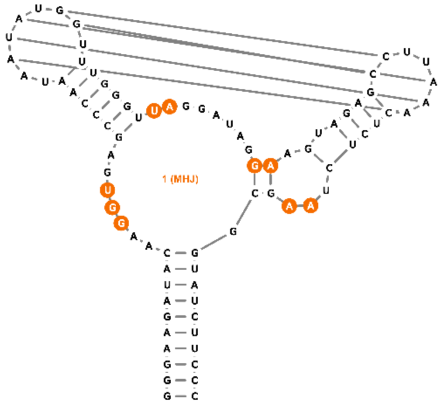

### Pseudoknots

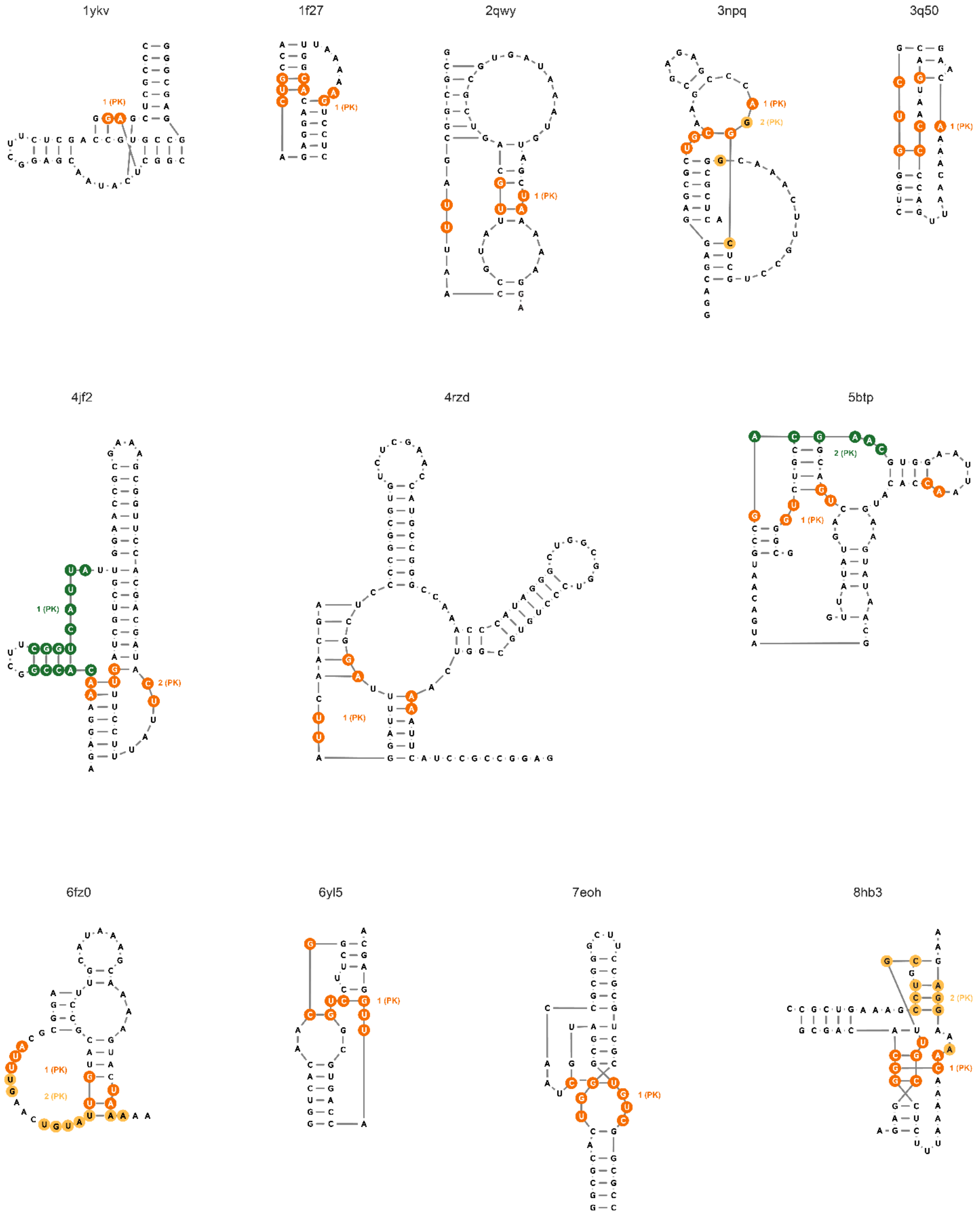

### G-quadruplexes

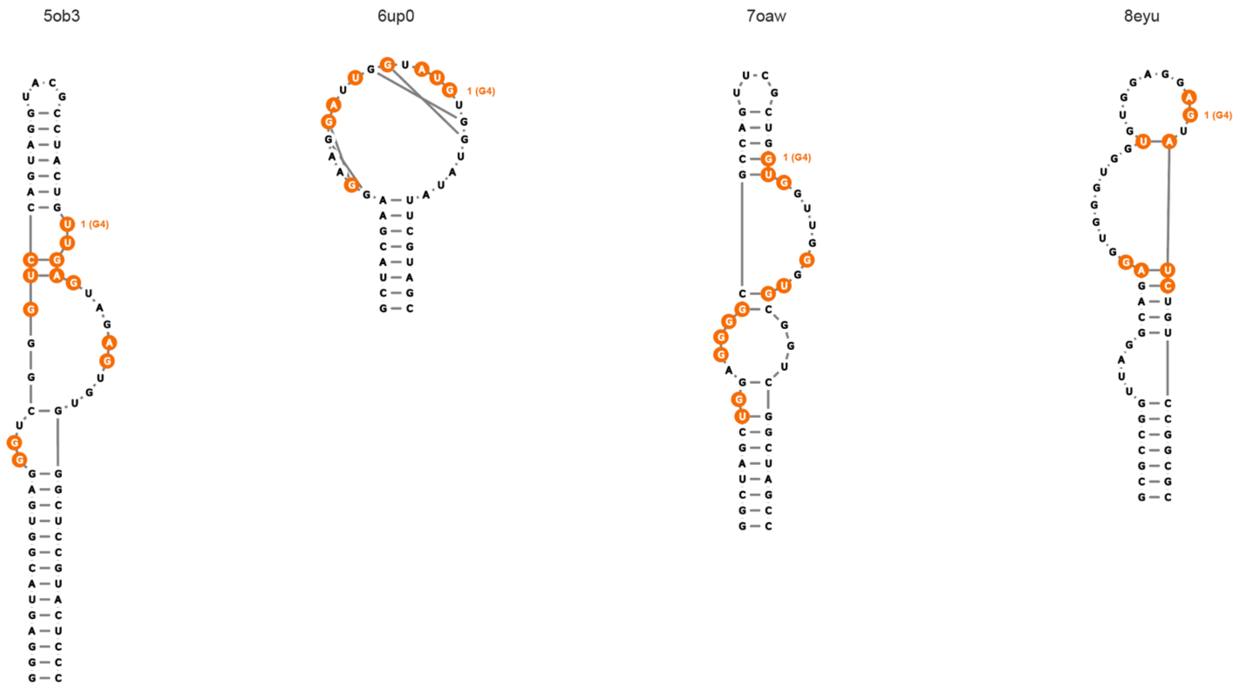


**Figure S6.** (5 panels, *above*) Secondary structural motifs that form pockets in our curated non-redundant RNA-ligand library, featuring ligands with high QED values. Pockets that overlap a known binding site of a drug-like ligand or an RNA–protein-residue contact in the structure are colored orange to yellow, the remaining pockets are considered novel and colored green to pink. Structures are classified as: (simple) simple structure, (con loop) consecutive loops, (MHJ) multi-helix junction, (PK) pseudoknot, or (dis) distant long-range interactions. PDB codes for all structures are provided in the Supporting Tables (below).

**16S rRNA Domain 5′** (7K00)

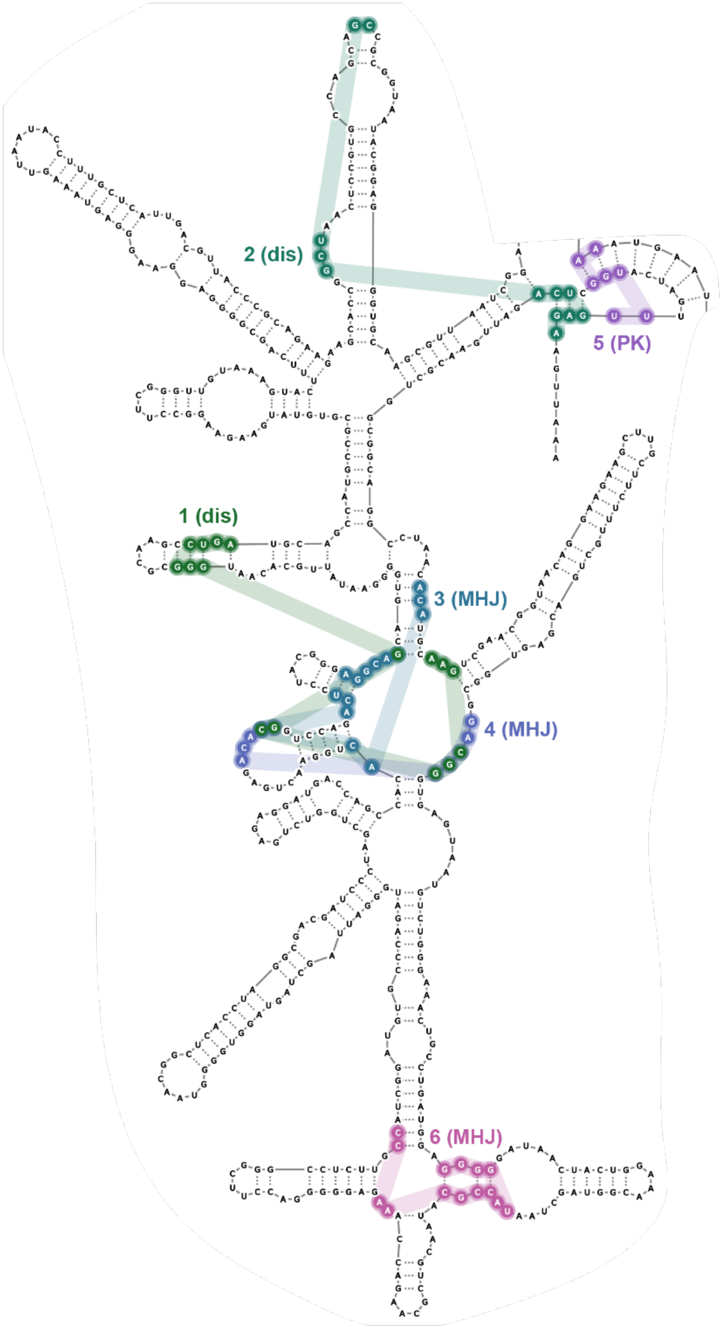

**16S rRNA Domain C** (7K00)

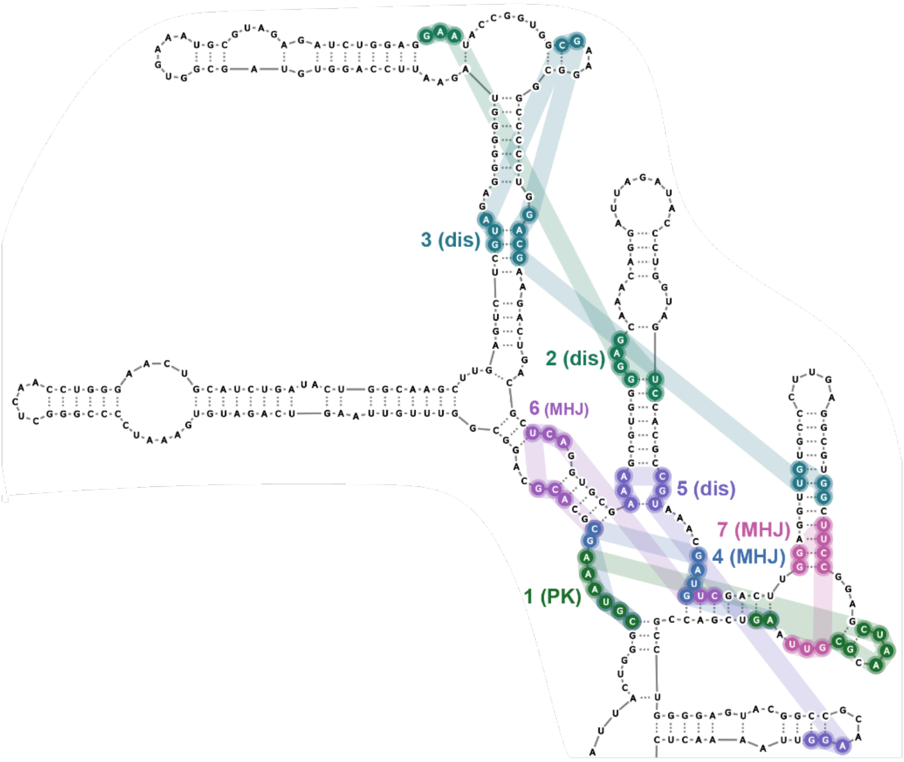

**16S rRNA Domain 3′M** (7K00)

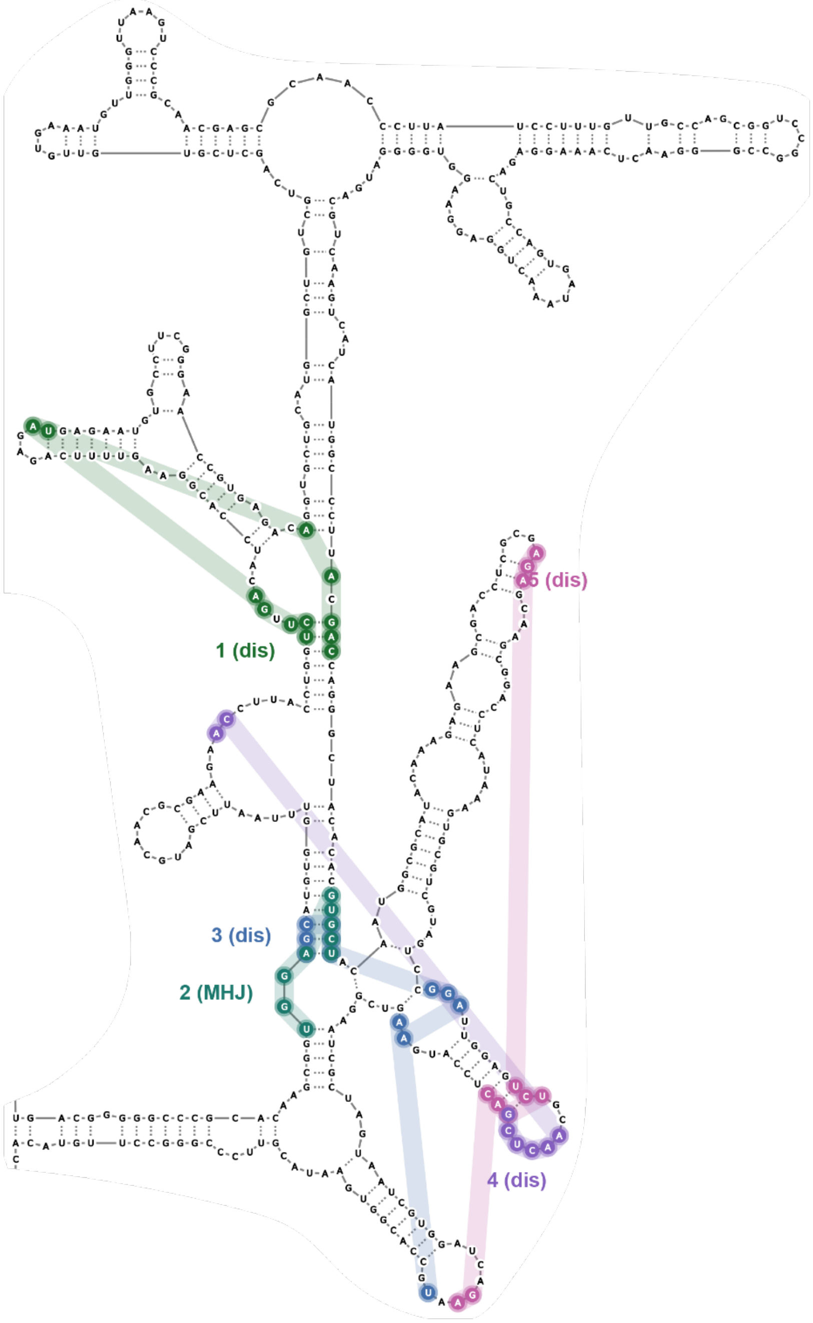

**16S rRNA Domain 3′m** (7K00)

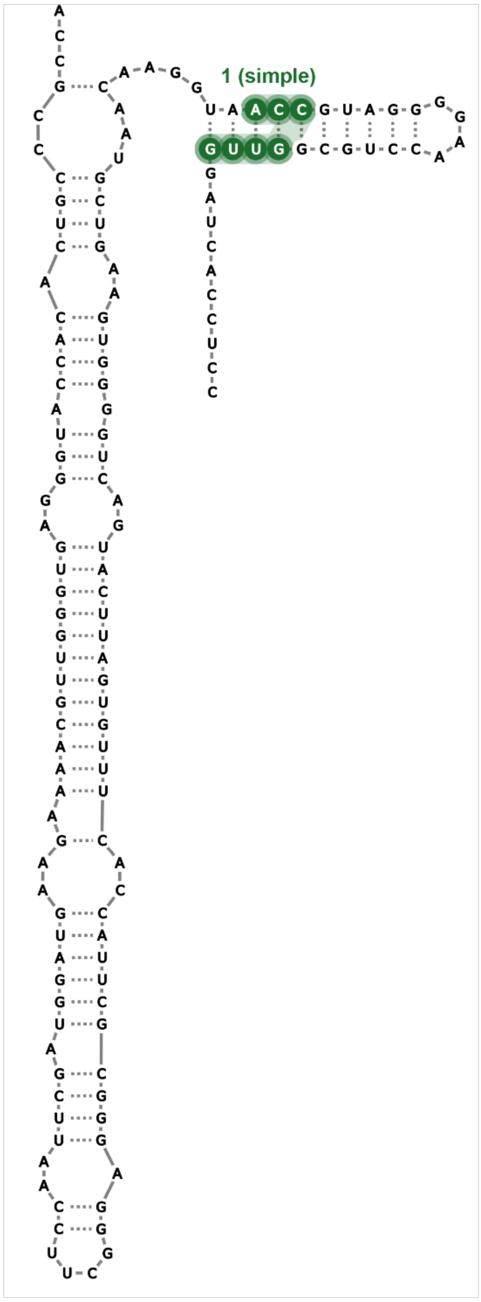

**23S rRNA Domain 0** (7K00)

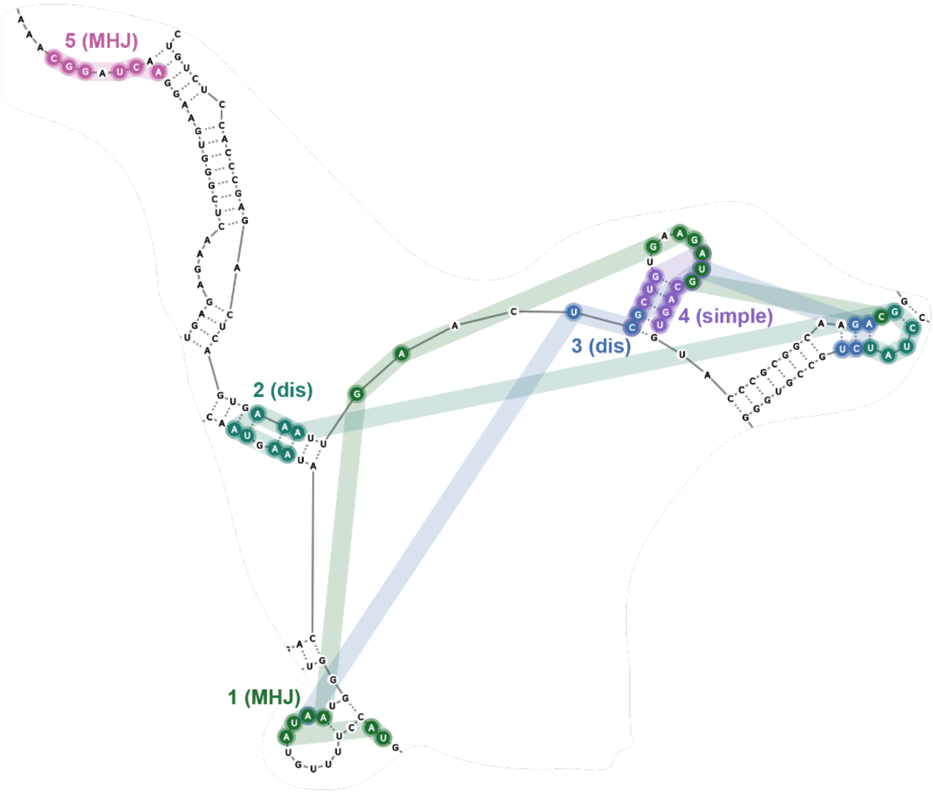

**23S rRNA Domain 1** (7K00)

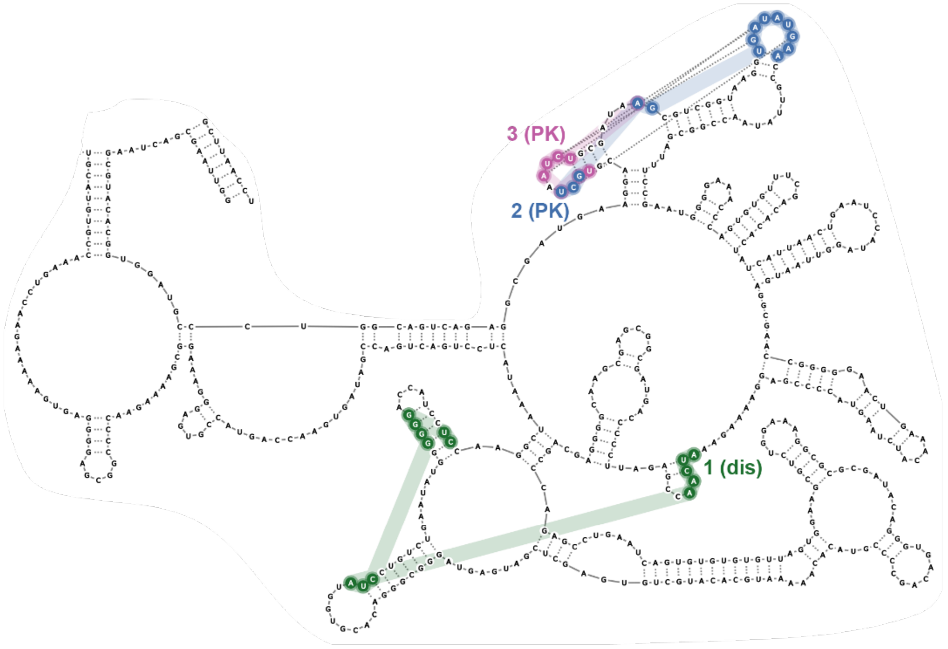

**23S rRNA Domain 2** (7K00)

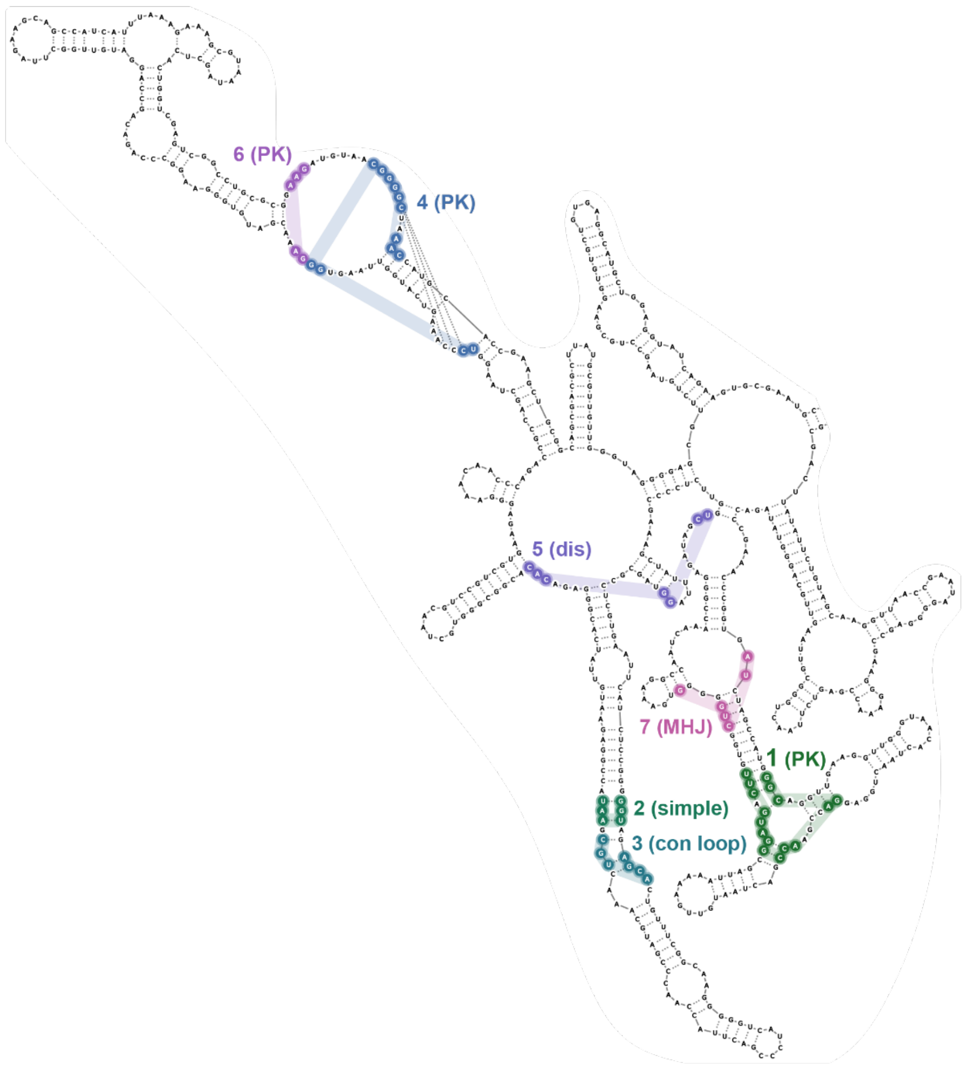

**23S rRNA Domain 3** (7K00)

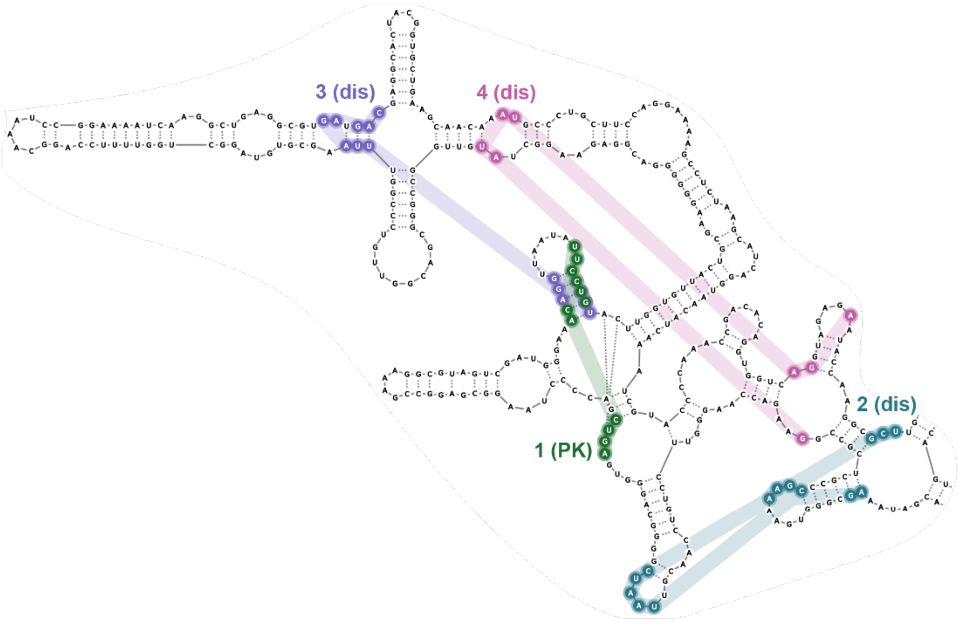

**23S rRNA Domain 4** (7K00)

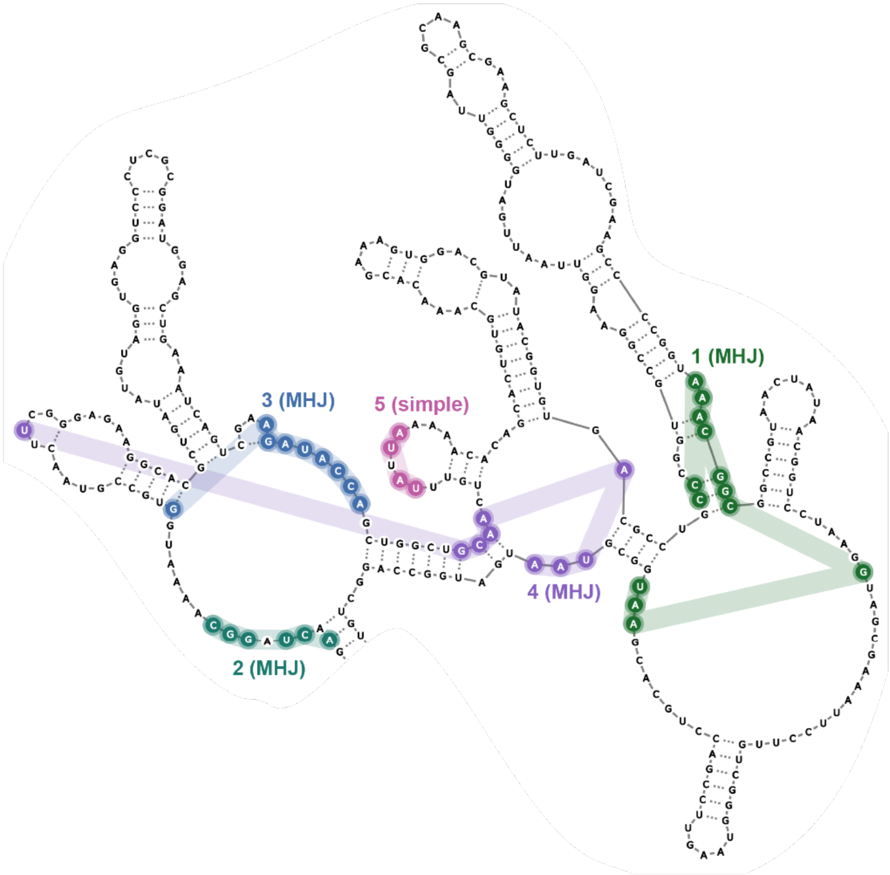

**23S rRNA Domain 5** (7K00)

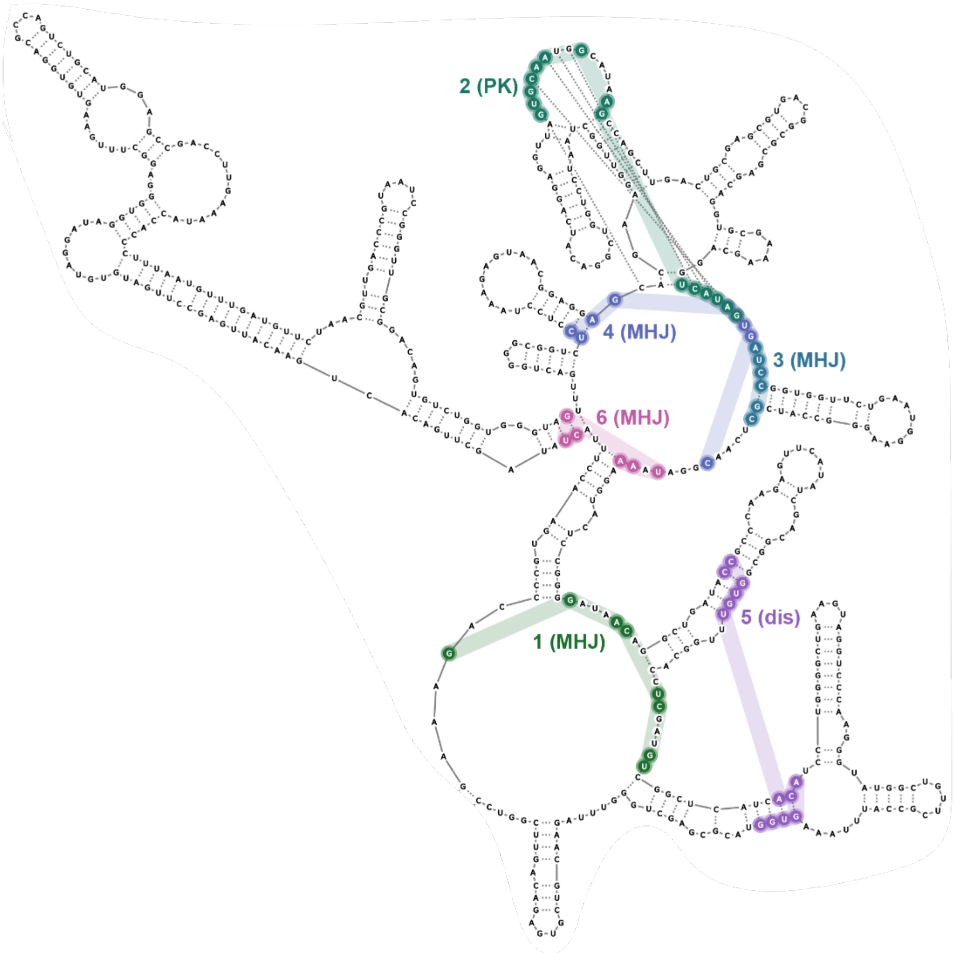

**23S rRNA Domain 6** (7K00)

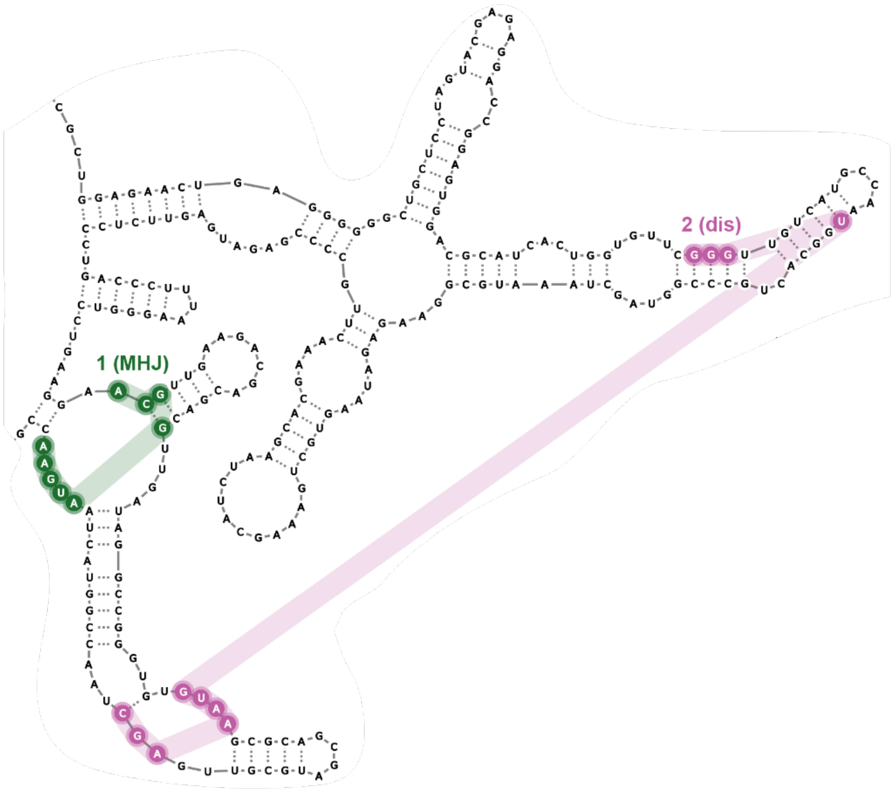


**Figure S7.** (11 panels, *above*) Secondary structural motifs that form pockets in each domain of the *E. coli* 16S and 23S rRNAs (PDB: 7K00) (12). Nucleotides that contact the same pocket are annotated with a number and matching color. Structures are classified as: (simple) simple structure, (con loop) consecutive loops, (MHJ) multi-helix junction, (PK) pseudoknot, or (dis) distant long-range interactions.

**Table S1.** Training set library.

| PDB code | reference | length | ligand name | QED |
|---|---|---|---|---|
| 1F27 | (19) | 30 | Biotin | 0.49 |
| 8D2B | (20) | 33 | TAL2 | 0.83 |
| 1Q8N | (21) | 38 | Malachite green | 0.75 |
| 7ELR | (22) | 45 | Xanthine | 0.45 |
| 3E5C | (11) | 53 | SAM (III) | 0.34 |
| 6FZ0 | (23) | 53 | SAM (V) | 0.34 |
| 3NPQ | (24) | 54 | SAH | 0.35 |
| 6UBU | (25) | 67 | Guanine | 0.46 |
| 2GDI | (26) | 80 | TPP | 0.79 |
| 6LAS | (27) | 93 | SAM (VI) | 0.34 |
| 4B5R | (28) | 94 | SAM (I) | 0.34 |
| 4RZD | (29) | 101 | PreQ1 (III) | 0.46 |
| 5KX9 | (30) | 112 | FMN | 0.33 |
| **Average** | | **66** | **-** | **0.48** |

**Table S2.** Test set library.

| PDB code | reference | length | ligand name | QED |
|---|---|---|---|---|
| 7FJ0 | (31) | 20 | KG022 | 0.63 |
| 6VA4 | (32) | 21 | MIP | 0.70 |
| 1LVJ | (33) | 31 | Acetylpromazine | 0.76 |
| 3Q50 | (34) | 33 | Pre-Q1 (I) | 0.46 |
| 6YL5 | (35) | 35 | SAH (SAM-SAH) | 0.35 |
| 2KTZ | (36) | 38 | Isis-11 | 0.87 |
| 6UP0 | (37) | 38 | YO3-biotin | 0.66 |
| 6XB7 | (38) | 41 | DMA-135 | 0.40 |
| 6GZR | (39) | 48 | 5-TAMRA | 0.38 |
| 1YKV | (40) | 49 | Ethanoanthracene | 0.64 |
| 7EOH | (41) | 49 | HBC | 0.68 |
| 8EYU | (42) | 49 | DFAME | 0.68 |
| 2QWY | (43) | 52 | SAM (II) | 0.34 |
| 7OAW | (44) | 52 | DMHBI+ | 0.62 |
| 8HB3 | (45) | 55 | Nicotinamide riboside | 0.45 |
| 6XJQ | (46) | 58 | 2,3-disubstituted epoxide | 0.38 |
| 3SKI | (47) | 68 | 2'-Deoxy-guanosine | 0.51 |
| 5OB3 | (48) | 69 | DFHBI | 0.86 |
| 5KPY | (49) | 71 | 5-Hydroxytryptophan | 0.62 |
| 4LX5 | (50) | 71 | PPDA | 0.55 |
| 5BTP | (51) | 75 | ZMP | 0.31 |
| 4JF2 | (52) | 77 | Pre-Q1 (II) | 0.46 |
| 7KVT | (53) | 83 | DFHBI-1T | 0.67 |
| 7DWH | (54) | 102 | SAM | 0.34 |
| 3D0U | (55) | 161 | Lysine | 0.46 |
| **Average** | | **58** | **-** | **0.55** |

**Table S3.** Parameter combinations tested during 3 rounds of *fpocketR* optimization**.**

| Round | *m* | *M* | *i* | *D* |
|---|---|---|---|---|
| **Default** | **3.4** | **6.2** | **15** | **2.4** |
| **1** | 2.6 3.0 3.4 3.8 4.2 | 5.4 5.8 6.2 6.6 7.0 | 10 15 20 25 30 | 1.6 2.0 2.4 2.8 3.2 |
| **2** | 2.6 2.8 3.0 3.2 3.4 | 5.0 5.2 5.4 5.6 5.8 | 20 25 30 35 40 | 1.6 1.8 2.0 2.2 2.4 |
| **3** | 2.8 2.9 3.0 3.1 3.2 | 5.4 5.5 5.6 5.7 5.8 | 36 38 40 42 44 | 1.5 1.55 1.6 1.65 1.7 |
| **Optimized** | **3.0** | **5.7** | **42** | **1.65** |

Note: Optimal parameters from each round are highlighted in orange.

**Table S4.** Summary of pocket detection performance in small RNA dataset.

| parameters | avg # of pockets | sens | rank 1 sens | ppv |
|---|---|---|---|---|
| ***fpocket*** *(protein optimized)* | 4.7 | 87% | 63% | 19% |
| ***fpocket-R*** *(RNA optimized)* | 1.3 | 100% | 92% | 78% |

**Table S5.** Detection of ligand binding pockets in paired apo and holo RNA structures.

| name | QED | state | PDB code | reference | resolution (Å) | known binding site |
|---|---|---|---|---|---|---|
| linezolid | 0.89 | apo | 7k00 | (12) | 1.98 | detected |
| | | holo | 3cpw | (16) | 2.70 | detected |
| FMN | 0.72 | apo | 6wjr | (56) | 2.70 | - |
| | | holo | 5kx9 | (30) | 2.90 | detected |
| adenine | 0.53 | apo | 5e54 | (57) | 2.30 | detected |
| | | holo | 5swe | (57) | 3.00 | detected |
| pre-Q1 | 0.46 | apo | 6vuh | (58) | 2.00 | detected |
| | | holo | 6vui | (58) | 2.98 | detected |
| lysine | 0.43 | apo | 3d0x | (55) | 2.95 | detected |
| | | holo | 3d0u | (55) | 2.80 | detected |
| TPP | 0.35 | apo | 8f4o | (59) | 3.10 | detected |
| | | holo | 2gdi | (26) | 2.05 | detected |
| SAM-I | 0.34 | apo | 3iqp | (60) | 2.90 | detected |
| | | holo | 3iqn | (60) | 2.70 | detected |
| glmS | 0.29 | apo | 2gcs | (61) | 2.10 | detected |
| | | holo | 2z74 | (62) | 2.20 | detected |
| self-alkylating epoxide | 0.25 | apo | 6xjz | (46) | 2.49 | detected |
| | | holo | 6xjq | (46) | 1.71 | detected |
| PRPP | 0.23 | apo | 6dnr | (63) | 2.90 | detected |
| | | holo | 6ck5 | (64) | 2.49 | - |

**Table S6.** Pocket characteristics for small RNA library.

| PDB code | pocket | structure type | structure class | occupied |
|---|---|---|---|---|
| 1F27 | 1 | local | PK | biotin |
| 1LVJ | 1 | local | simple | acetylpromazine |
| 1Q8N | 1 | local | con. loops | malachite green |
| 1YKV | 1 | local | PK | ethanoanthracene |
| 2GDI | 1 | local | MHJ | TPP |
| 2KTZ | 1 | local | simple | Isis-11 |
| 2QWY | 1 | local | PK | SAM |
| 3D0U | 1 | local | MHJ | lysine |
| 3E5C | 1 | local | con. loops | Sr |
| 3E5C | 2 | local | MHJ | SAM |
| 3NPQ | 1 | local | PK | SAH |
| 3NPQ | 2 | local | PK | SAH |
| 3Q50 | 1 | local | PK | PreQ1 |
| 3SKI | 1 | local | MHJ | 2'-deoxy-guanosine |
| 3SKI | 2 | local | MHJ | - |
| 3SKI | 3 | local | MHJ | - |
| 4B5R | 1 | local | PK | Ba |
| 4B5R | 2 | local | MHJ | - |
| 4B5R | 3 | local | MHJ | SAM |
| 4JF2 | 1 | local | PK | Cs |
| 4JF2 | 2 | local | PK | PreQ1 |
| 4LX5 | 1 | local | MHJ | PPDA |
| 4RZD | 1 | local | PK | PreQ1 |
| 5BTP | 1 | local | PK | ZMP |
| 5BTP | 2 | local | PK | RNA (dimer) |
| 5KPY | 1 | local | MHJ | 5-hydroxy-L-tryptophan |
| 5KX9 | 1 | local | MHJ | ribocil-D |
| 5KX9 | 2 | local | PK | ribocil-D |
| 5OB3 | 1 | local | g-quadraplex | DFHBI |
| 6FZ0 | 1 | local | PK | SAM |
| 6FZ0 | 2 | local | PK | SAM |
| 6GZR | 1 | local | MHJ | 5-TAMRA |
| 6LAS | 1 | local | MHJ | SAM |
| 6UBU | 1 | local | MHJ | guanine |
| 6UP0 | 1 | local | g-quadraplex | YO3-biotin |
| 6VA4 | 1 | local | simple | MIP |
| 6XB7 | 1 | local | con. loops | DMA-135 |
| 6XJQ | 1 | local | con. loops | 2,3-disubstituted epoxide |
| 6XJQ | 2 | local | con. loops | protein |
| 6YL5 | 1 | local | PK | SAH |
| 7DWH | 1 | local | con. loops | SAM |

| | | | | |
|---|---|---|---|---|
| 7ELR | 1 | local | con. loops | xanthine |
| 7EOH | 1 | local | PK | HBC |
| 7FJ0 | 1 | local | simple | KG022 |
| 7KVT | 1 | local | MHJ | DFHBI-1T |
| 7OAW | 1 | local | g-quadraplex | DMHBI+ |
| 8D2B | 1 | local | con. loops | TAL2 |
| 8EYU | 1 | local | g-quadraplex | DFAME |
| 8HB3 | 1 | local | PK | NNR |
| 8HB3 | 2 | local | PK | NNR |

**Table S7.** Pocket characteristics for the *E. coli* 16S rRNA (PDB: 7K00) (12).

| domains | pocket | type | structure class | occupied |
|---|---|---|---|---|
| 1 | 1 | long-range | long-range | - |
| | 2 | long-range | long-range | - |
| | 3 | local | MHJ | - |
| | 4 | local | MHJ | - |
| | 5 | local | PK | - |
| | 6 | local | MHJ | Mg |
| | 7 | local | con. loops | Mg |
| | 8 | local | PK | - |
| | 9 | local | simple | - |
| | 10 | long-range | long-range | - |
| | 11 | local | MHJ | Mg |
| | 12 | long-range | long-range | - |
| 2 | 1 | local | PK | - |
| | 2 | long-range | long-range | - |
| | 3 | long-range | long-range | - |
| | 4 | local | MHJ | - |
| | 5 | long-range | long-range | - |
| | 6 | local | MHJ | - |
| | 7 | local | MHJ | - |
| | 8 | local | MHJ | - |
| | 9 | local | MHJ | protein |
| | 10 | local | simple | - |
| 3 | 1 | long-range | long-range | - |
| | 2 | local | MHJ | - |
| | 3 | long-range | long-range | - |
| | 4 | long-range | long-range | - |
| | 5 | long-range | long-range | - |
| | 6 | long-range | long-range | protein |
| | 7 | local | MHJ | Mg |
| | 8 | long-range | long-range | - |
| | 9 | long-range | long-range | - |
| | 10 | long-range | long-range | - |
| | 11 | long-range | long-range | - |
| | 12 | local | MHJ | Mg |
| 4 | 1 | local | simple | |

**Table S8.** Pocket characteristics for the *E. coli* 23S rRNA (PDB: 7K00) (12).

| domain | pocket | type | structure class | occupied |
|---|---|---|---|---|
| 0 | 1 | local | MHJ | RNA |
| | 2 | long-range | long-range | protein |
| | 3 | long-range | long-range | - |
| | 4 | local | simple | protein |
| | 5 | local | MHJ | RNA |
| | 6 | long-range | long-range | protein |
| | 7 | long-range | long-range | RNA |
| 1 | 1 | long-range | long-range | - |
| | 2 | local | PK | - |
| | 3 | local | PK | - |
| | 4 | local | MHJ | protein |
| | 5 | local | simple | RNA |
| 2 | 1 | local | MHJ | - |
| | 2 | local | simple | RNA |
| | 3 | local | Con. Loops | - |
| | 4 | local | PK | - |
| | 5 | long-range | long-range | RNA |
| | 6 | local | PK | - |
| | 7 | local | MHJ | - |
| | 8 | local | simple | RNA |
| 3 | 1 | local | PK | - |
| | 2 | long-range | long-range | - |
| | 3 | long-range | long-range | - |
| | 4 | long-range | long-range | - |
| | 5 | local | MHJ | - |
| | 6 | local | MHJ | - |
| | 7 | local | MHJ | - |
| 4 | 1 | local | MHJ | - |
| | 2 | local | MHJ | - |
| | 3 | local | MHJ | - |
| | 4 | local | MHJ | - |
| | 5 | local | simple | RNA |
| 5 | 1 | local | MHJ | linezolid |
| | 2 | local | PK | - |
| | 3 | local | MHJ | protein |
| | 4 | local | MHJ | - |
| | 5 | long-range | long-range | RNA |
| | 6 | local | MHJ | RNA |
| | 7 | long-range | MHJ | RNA |
| | 8 | local | MHJ | - |
| 6 | 1 | local | MHJ | - |

| | | | | |
|---|---|---|---|---|
| | 2 | long-range | long-range | - |

**Table S9.** Pocket characteristics for the group II intron (PDB: 5G2X) (65).

| pocket | type | structure class | occupied |
|---|---|---|---|
| 1 | local | MHJ | - |
| 2 | local | MHJ | - |
| 3 | long-range | long-range | - |
| 4 | local | MHJ | - |
| 5 | local | MHJ | - |
| 6 | long-range | long-range | - |
| 7 | local | MHJ | - |
| 8 | long-range | long-range | - |
| 9 | local | simple | - |
| 10 | local | simple | - |
| 11 | long-range | long-range | - |

**Table S10.** Summary of pocket characteristics for protein-ligand versus RNA-ligand complexes.

| library | parameters | pocket type | score | # alpha spheres | SASA | volume | hydro-phobic density | hydro-phobicity score | polarity score |
|---|---|---|---|---|---|---|---|---|---|
| **RNA** (n=38) | **fpocket-R** (RNA optimized) | **all** | 0.34 | 84 | 117 | 351 | 8.17 | -1.1 | 4.6 |
| | | **rank 1** | 0.38 | 93 | 119 | 369 | 9.85 | -1.0 | 4.6 |
| | | **known** | 0.36 | 90 | 116 | 361 | 9.45 | -1.8 | 4.5 |
| | **fpocket** (protein optimized) | **all** | 0.02 | 38 | 134 | 472 | 1.72 | 1.2 | 4.5 |
| | | **rank 1** | 0.23 | 61 | 146 | 560 | 4.92 | -0.8 | 5.2 |
| | | **known** | 0.17 | 96 | 206 | 750 | 7.79 | -1.8 | 6.3 |
| **protein** (n=200) | **fpocket-R** (RNA optimized) | **all** | 0.35 | 64 | 95 | 318 | 19.89 | 24.2 | 7.7 |
| | | **rank 1** | 0.49 | 77 | 103 | 370 | 21.12 | 25.6 | 8.8 |
| | | **known** | 0.38 | 77 | 119 | 401 | 19.10 | 22.7 | 9.2 |
| | **fpocket** (protein optimized) | **all** | 0.03 | 32 | 103 | 342 | 9.59 | 18.1 | 5.7 |
| | | **rank 1** | 0.28 | 83 | 169 | 635 | 19.81 | 26.6 | 10.6 |
| | | **known** | 0.16 | 84 | 191 | 676 | 21.10 | 24.6 | 10.8 |
